# Quantifying the ∼75-95% of Peptides in DIA-MS Datasets that were not Previously Quantified

**DOI:** 10.1101/2024.12.15.628560

**Authors:** Gautam Saxena, Qin Fu, Aleksandra Binek, Jennifer E Van Eyk

## Abstract

We demonstrate an algorithm termed GoldenHaystack (GH) that, compared to the leading DIA-MS algorithm, (a) quantifies and identifies with better FDR accuracy the peptides found in FASTA search spaces (∼5-25% of analytes in DIA-MS datasets), (b) quantifies the remaining ∼75-95% of analytes that were previously unquantified, and (c) runs ∼40-200x faster (or ∼1-10x faster than the LC-MS). Specifically, without a FASTA or spectral library, GH can deconvolute and accurately quantify chimeric LC-MS spectra. The central idea that enables this claim is: for sufficiently sized projects (e.g., ≥ ∼50 LC-MS files), pairs of peptides that co-elute in one subset of LC-MS files do not exactly co-elute in a different subset of files. GH thus analyzes a project holistically: it uses *multi*-partite matching to match fragment ions across all samples, separates and regroups the fragment ions into unique analyte signatures, reduces stochastic noise, and then quantifies those unique analyte signatures.

## I. Introduction

Data independent acquisition^1^ (DIA) liquid chromatography mass spectrometry (LC-MS) proposes to record all the MS2 fragment information for all peptides in the LC-MS dataset instead of first preselecting a comparatively small set of peptides at the MS1 stage and only subsequently performing MS2 fragmentation on this subset, as done in data-dependent acquisition. However, the benefits from the DIA-MS protocol are achieved at the cost of producing highly chimeric spectra: for any given liquid chromatography (LC) elution period, the set of ion chromatogram (XIC) traces is frequently a composite of two or more peptides’ MS2 fragment XIC traces (Fig. 1a).

**Fig. 1:**
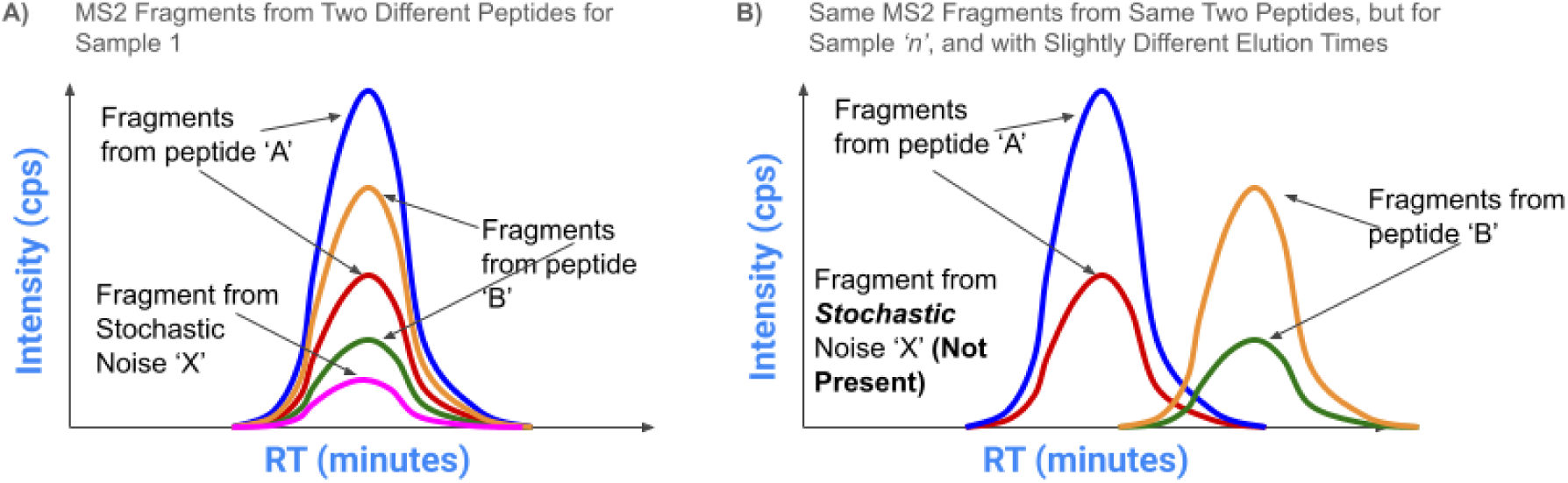
Co-elution of Two Peptides. **A**) Simplified illustration of 5 MS2 fragments from two different but co-eluting peptides (2 MS2 fragments from peptide ‘A’ and another two from peptide ‘B’, plus one fragment from a co-eluting stochastic noise analyte). Because the MS2 fragments are co-eluting, one cannot determine from Fig. 1A alone whether all five fragments belong to a single peptide, multiple peptides, noise analytes, or some combination thereof. **B**) The same 2 peptides elute in sample *‘n’*, but now at slightly different elution (RT) times. Both visually and computationally, one can then determine that the four MS2 fragments correspond to at least two distinct peptides. Moreover, the fragment resulting from the stochastic noise analyte is no longer present (i.e., it is “stochastic”), and so it too can be computationally removed from inclusion when generating unique analyte signatures. Finally, after proper deconvolution, a computational algorithm can ensure that the MS2 fragments corresponding to the green and yellow XIC traces not be used to quantify peptide ‘A’, and similarly the MS2 fragments corresponding to the blue and red XIC traces not be used to quantify peptide ‘B’.

To accurately quantify the peptides present in these chimeric LC-MS spectra, the last decade of DIA algorithms (e.g., DIA-NN^2^, Spectronaut^3^, DIA-umpire^4^, OpenSwath^5^ etc.) used peptide sequence information derived from protein FASTA files and, to ensure a controlled search space, allowed users to preselect only a very small set of variable PTMs (typically 1 or 2) for the DIA search algorithms to consider. However, as recent analysis has shown that these genomics-derived FASTA files are substantially incomplete^6,7^ and there are hundreds of possible PTMs^8^, existing DIA algorithms are not quantifying a large proportion of analytes present in DIA-MS files. Alternative methods, such as de novo sequencing of DIA datasets^9,10^, have been proposed over the years as an alternative to help address the issue of incomplete identifications from FASTA files, but the de novo DIA-MS algorithms tend to take substantially longer than even FASTA-based algorithms, have limited support for PTMs, and identify substantially fewer FASTA-based peptides than the leading DIA-MS (FASTA-based) algorithms (Note S1).

More pointedly, unexplored PTMs (e.g., glycosylation, citrullination, ubiquitination, acetylation, methylation, phosphorylation, sumoylation, and hundreds more) and unpredicted peptide sequences (e.g., splice variants, sequence variation, protein precursors, unusual proteolytic cleavages, novel proteins, etc.) are more likely to be disease-specific^11^ or disproportionally biologically influential^7,12^ and thus may be beneficial in separating study conditions. An example that suggests the importance of unpredicted peptide sequences and unexplored PTMs is the very recent biomarker study on Alzheimer^13^, in which the two possible biomarkers the authors proposed are related to (a) the abundance of a form of the protein amyloid-β in which there were two extra amino acids at its C-terminal end (which would result in a non-standard peptide sequence in the MS) and (b) the presence of a PTM, specifically a phosphate group, on the 217^th^ amino acid of the protein tau instead of the non-phosphorylated form of tau. While phosphorylation may be a PTM that some researchers preselect for consideration in DIA experiments, there are hundreds of other biologically important PTMs^8^ that may differentiate one study condition from another. Similarly, while some FASTA files can contain non-standard sequences (e.g., isoforms) in what is referred to as non-canonical FASTA files, (a) they take often far longer to computationally process, (b) the FDR rates explode with the much larger search space, and (c) importantly, even these non-canonical FASTA files may still miss disease-specific protein sequences, such as unexpected proteolytic cleavages or new proteins not predicted by traditional genomics sequencing algorithms (e.g., microproteins^7^, such as the microprotein ASNSD1-uORF^12^ that is implicated in medulloblastoma tumors, a brain cancer that affects children). Finally, we recently learned that the Mann lab had published a paper as far back as early 2011 in which they showed that while ∼10k spectra were identified in a DDA experiment of a HeLa cell lysate with long gradients (i.e., up to ∼250 min), over 100k peptides were detected but not identified^14^. Interestingly, those approximate stats (a rough 10-to-1 ratio between detectable peptides vs identifiable ones) are similar to what we show for GH (Table 6, dataset #1, 2, and 4), except that we are (a) using DIA, (b) typically much shorter gradients (∼25min), (c) human plasma instead of cells (for datasets #1, 2 and 4), and (d) we quantify all those unidentified analytes and show that their linearity response for quantitation is similar to those of identified analytes.

Although deconvoluting and quantifying all the spectra in DIA-MS datasets without using FASTA files or spectral libraries can be very valuable, if such a bioinformatics algorithm were to take too long to run, it would be effectively unusable for data analysis. Currently, the leading DIA-MS algorithms (which quantify only the FASTA identifiable peptides) either take too long to run (i.e., often >> 2x longer than the MS acquisition time for non-Astral datasets) or, for the larger data sets with the Astral MS, may not complete in any reasonable timeframe when as few as three variable modifications are considered. For example, on a 61 sample project which was run on an Exploris 480 and searched with a combination of MSFragger and DIA-NN with three variable modifications considered (oxidation on M, phosphorylation on S/T/Y, and stable isotopic label on K/R), the MSFragger-DIA-NN based informatics portion took, on a cluster, 90.72 hours (ignoring the “load conditions” described in Fig. 5) or ∼500 hours (including the load conditions), whereas the LC-MS acquisition time was ∼24 hours. And for an Astral dataset, which had file sizes that were ∼5x-10x larger than the Exploris 480 file sizes (i.e., ∼8GB / file), MSFragger + DIA-NN, when including those same three variable modifications, could not be run in any reasonable timeframe. In short, given modern LC-MSs’ current file sizes coupled to the increasing number of projects in which the number of LC-MS files per project exceeds ∼50, the leading DIA-MS informatics algorithms have runtimes that constrain and dictate what biological questions can be asked (e.g., users are recommended to not search more than one variable modification), even though the DIA-MS datasets may contain the answers.

The goal is to have a deconvolution strategy that can accurately quantify the majority of peptides present in a DIA-MS datasets without using – and being constrained by – FASTA files, spectral libraries, or user’s preselected PTMs, and to do so quickly. That is, the informatics pipeline should ideally be as unbiased and at least as fast as the LC-MS instruments, but preferably much faster.

## II. Computational Methods

Although we ultimately use a different technique to deconvolve chimeric LC-MS spectra, we first share the original technique, as it simple and illustrative: instead of the LC-MS running *‘n’* samples with the same identical starting-stopping boundaries for the MS2 DIA windows for each sample (“identical windows”), we run the *‘n’* samples with *‘n’* slightly different starting-stopping boundaries for the MS2 windows (“*n*-Offset Windows”). For example, sample 1’s MS2 DIA windows could be set at …625-650, 650-675, 675-700… while sample 2’s could be slightly shifted to 637.5-662.5, 662.5-687.5, 687.5-712.5… and sample 3’s could be shifted once more to …643.7-668.7, 668.7-693.7, 693.7-718.7… and so on for all ‘n’ samples (Fig. 2). This protocol may appear similar to the overlapping window approach by the MacCoss lab^15^, but it is instead very different in both technique and benefits, as discussed further in Note S2. With sufficient samples analyzed, we can then computationally determine through mathematical set operations which peptide fragment ions belong together to form unique analyte signatures (UAQs) without resorting to any pre-existing FASTA file or spectral library.

**Fig. 2:**
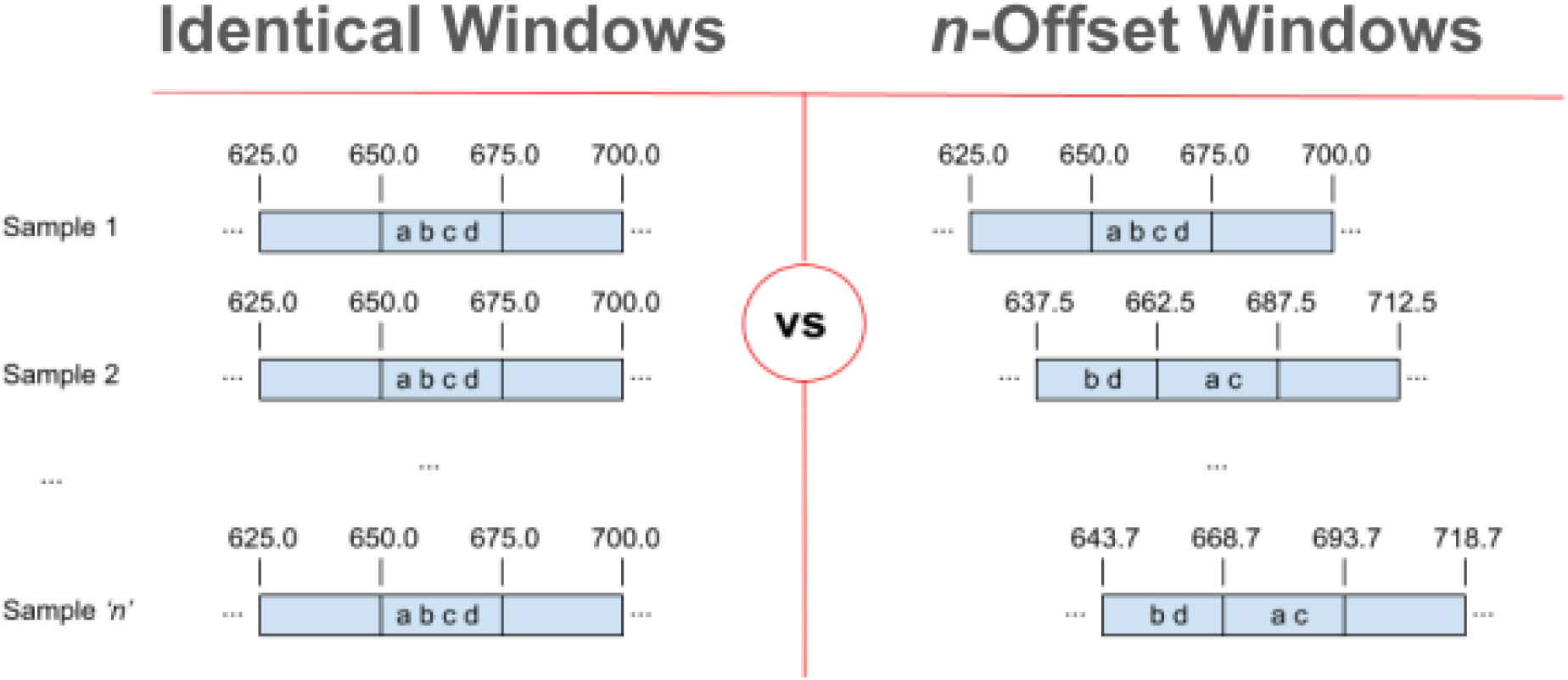
Slightly Different DIA Window Ranges for Different Samples. The left panel, named “Identical Windows”, is a simplified illustration of the standard DIA protocol: for a project of *‘n’* samples, all samples have the exact same set of DIA MS2 windows, with exact starting / ending values. Consequently, for the four MS2 fragments represented by the letters ‘a’, ‘b’, ‘c’, and ‘d’, one cannot determine if those four MS2 fragments belong to a single peptide or multiple peptides, i.e., one cannot computationally separate the m/z fragments given the available information. In the right panel, however, the ‘n’ samples have ‘n’ slightly different starting / stopping DIA windows, but still of identical widths. Consequently, in this illustration, the four fragments do not always appear in the same DIA window bucket. One can therefore computationally determine that a) the four fragments belong to two or more peptides and, as a bonus, b) the precursor m/z corresponding to fragments ‘b’ and ‘d’ can be narrowed from the original 25Th range (e.g., 650 to 675Th) to a narrower range of 650 to 662.5Th (a 12.5Th range) in this highly simplified 3 sample illustration. For ‘n’ samples, the theoretical max narrowing of precursor m/z range and in-silico separation abilities is linearly proportional to *‘n’* (i.e., the range is reduced to 1/n * original window width), but in practice it is limited by physical constraints – e.g., sharpness of MS quadrupole filter – to ∼0.07Th.

However, although this “*n*-Offset Window” approach appeared to work in our initial tests, it had a number of significant drawbacks: (a) it was far too computationally intensive, despite many advanced optimizations and availability of thousands of CPUs; (b) it would have required MS labs to re-do their experiments using this new protocol; and, (c) existing DIA software, such as DIA-NN, may not have functioned properly against this unexpected offsetting of the MS2 windows for every sample.

Instead, we took advantage of the *natural* variation in LC elution times for any given pair of peptides^16^, which provided similar benefit to the “*n*-Offset Window” approach described above; i.e., since peptides that exactly co-elute in one subset of MS samples may not exactly co-elute in other subset of MS samples (Fig. 1b), one can computationally separate which MS2 fragments belong as a single set (*i.e.,* a single peptide) compared to two or more different sets (i.e., two or more different peptides). To achieve this in-silico separation, we employed a weighted multi-partite matching algorithm based on the relatively recent and efficient algorithm^17^ by Da Tang and Tony Jebara, director of Netflix’s Artificial Intelligence (AI) / Machine Learning (ML) unit, that used coordinate descent to arrive at an approximate matching solution within a reasonable timeframe. When using this multipartite matching algorithm, we performed the matching of MS2 XIC fragments across all of a project’s samples using the MS2 XIC’s mass-to-charge (m/z) and retention time (RT) similarities. At a very high level, this matching may appear similar to the standard “Match Between Runs” (MBR) logic commonly used in DDA algorithms such as MaxQuant^18^ and IonQuant^19^ or to pure feature alignment algorithms such as the one popularized by Progenesis QI^20^. However, this multi-partite matching is distinct in at least three fundamental ways: (a) first and foremost, in contrast to MBR, the matching occurs for *all* MS2 fragment ions irrespective of whether the fragment ions have been identified in any sample from a FASTA search space. In MBR algorithms, the matching is limited to using only those peptides identifiable from FASTA search spaces (∼5-25% of peptides in DIA-MS datasets). (b) Secondly, in contrast to pure feature alignment algorithms, GH barely uses MS1 information and relies almost exclusively on MS2 fragment information, whereas pure feature alignment programs exclusively use the MS1 information for alignment, i.e., they do not use MS2 information for feature alignment. However, in DIA experiments, MS1 signals are often non existent even though the corresponding MS2 fragment ions are clearly present, and when the MS1 signals do exist, they are almost always far too convolved to be accurately used in generating feature alignment in complex tissues such as human plasma or cell tissue, and this issue becomes more pronounced the more the gradient length is reduced (e.g., from ∼60 min to <30 min). (c) Third and lastly, GH’s matching employs a multipartite instead of a bipartite technique, which means that GH tries to optimize the matching of fragment ions across all samples simultaneously, instead of matching fragment ions from only two samples at a time in serial. Serial bipartite matching can cause errors from an early matching pair to propagate to subsequent sample pairs. Further, serial bipartite matching technique does not allow useful information from subsequent samples to inform the ideal matching between any previous two samples. In other words, multi-partite matching is akin to a large jigsaw puzzle where it is not sufficient for two pieces to more-or-less fit well together (i.e., bipartite match) – instead, all the pieces must be *collectively* optimized to fit well together (i.e., multipartite match) in order for any two pieces to be considered well matched, thus reducing overall fragment ion match errors. That said, for performance optimization reasons, after a certain threshold number of samples, e.g., 50, the full multi-partite matching described in the original “Netflix paper” is changed to a semi multi-partite matching algorithm in which the first ∼50 samples fragment ion matchings are considered “locked in” and can no longer change due to information from other samples. None of the drawbacks described in the previous paragraph on the “*n*-Offset Window” approach applied, and as an unexpected additional benefit, each MS2 DIA window could be processed in parallel since the data in each MS2 DIA window was completely independent of every other MS2 DIA window. Further, another unanticipated benefit was that this same deconvolution logic could automatically reduce stochastic noise, since the signals resulting from noise (e.g., fragments from random analytes present on the LC column, in the LC-MS room’s air, generated by electronic chatter etc.) were random, and rarely appeared in sufficient number of samples at the exact same co-eluted RT location. We have termed this entire process “Jitter Deconvolution”, owing to the natural, inherent variation – the jitter – in LCs for datasets with ≥ ∼50 files.

As an aside to this paper’s core computational method, we also describe an alternative and straightforward means for users to gain potential insight into any algorithm’s claimed FDR rates, as follows: when performing a search against a FASTA file, we add a variable modification on K (+8.0142Th) and R (+10.0083Th). We chose these particular variable modifications because they correspond to stable isotopically labelled (SIL) modifications that all search engines make easily available, typically from a drop-down list, and these mass shifts are not known to be present in nature, which may be one reason why they have been used for several decades in multiplex reaction monitoring (MRM) MS clinical validation settings. Once the search has completed, we then count the number of PSMs identified at the desired FDR threshold (e.g., 1% FDR) that have these SIL modifications. Assuming no synthetic peptides with SIL peptides were spiked into the samples, the true number of SIL PTMs is 0, but acceptable answers would be between 0 and approximately the number of PSMs corresponding to the user-selected FDR rate. (More details regarding SIL search requirements can be found in Note S3). This SIL-searching approach – proposed by Dr. Martin Frejno to be named “Within Organism Entrapment” (WOE) – can complement the traditional entrapment (TE) approach in which one searches a special concatenated FASTA file formed by merging the normal FASTA file (n-FASTA) with a different organism’s (e.g., *C. elgans*) FASTA file (d-FASTA). However, although WOE and TE are similar in both approach and purpose, there are three main differences: (a) the WOE approach does not require a bioinformaticist’s time, and a typical user can accomplish the WOE setup in less than ∼15 seconds of effort, which is in contrast to TE’s need of a bioinformaticist’s time of several hours to do TE accurately (Note S4); (b) with WOE, there is no debate or ambiguity when SIL peptides are identified that they are definitively false matches, since SIL modifications do not occur in nature; in contrast, in TE, when peptides from the d-FASTA file are identified, there is always the argument – which some of the authors on this paper have expressed in past experiments – that possibly those identified peptides from the different organism are not false matches but are real peptides since FASTA files are incomplete (e.g., these d-FASTA peptides could be legitimate splice variants, SNPs, proteolytic cleavages, precursors, novel proteins not predicted by genomics sequencing that should have existed in the n-FASTA file etc.). This scenario is particularly pronounced when the informatics algorithm claims a peptide-to-spectra match with high confidence and visual inspection of the spectra shows plausible matching to the claimed peptide sequence, as occurred in Table 5; and (c) there may be a frequency and grammar to how amino acids appear next to each which differ by organisms, just as with the English language the letter “i” typically follows “e” except after “c”, and not the other way around, but that childhood spelling rule may not apply in other Latin script languages. In the case of WOE, those hidden amino acid rules, if any, are fully preserved: the frequency and order of amino acids in peptides and the frequency histogram for the number of amino acids in a peptide (and even other attributes, such as the typical terminal amino acid in a peptide from the terminal portion of a protein) is identical between SIL peptides and non SIL peptides. With TE, none of these amino acid “grammar rules” may be similar in the d-FASTA file compared to the n-FASTA file, and so the informatics algorithms may be unfairly biased to these hidden amino acid rules, i.e., the informatics algorithms may have a bias to the hidden amino acid “grammer” in one organism vs another.

Finally, to achieve fast processing, we use (a) stream processing (i.e., data rarely is stored on disk, where access times can be slow, but instead the output of one step is immediately used as the input in the next step, which also helps substantially with CPU caching) and (b) massive single-server parallelism, not just across all CPU threads available in even the largest servers but even within a single CPU through use of vector processing, i.e., SIMD processing. (SIMD on chip processing is a very light-weight and far smaller version of GPU processing, but with faster data communication between the core CPU chip and the SIMD layer.) As well, we used 4^th^ Generation AMD EPYC processors which have 64 bit vector SIMD support as opposed to only 32 bit vector SIMD support as common in older CPU architectures. As an illustration of the challenge of designing an algorithm to be truly massively parallel, the leading DIA-MS algorithm of DIA-NN does parallelize processing, but it does not always use all the available cores, sometimes being limited to ∼32 cores even if hundreds are available^21^. Or, informatics algorithms may be limited for other reasons, such as possibly using slow disk access instead of fast RAM due to too much RAM pressure causing the operating systems to swap^22^. In contrast, GH achieves nearly linear performance with the number of threads, and the results shown in Fig. 5 were achieved on a 360 thread server with CPU utilization above 90% during the vast majority of the processing time. That is, we used most of the 360 cores most of the time and so should a new server with even more cores become available, GH’s performance would increase linearly with the increase in number of cores, without any further code customization to GH. As a final bonus, GH’s algorithm generate a single consolidated spectra file for FASTA searching, irrespective of the number of LC-MS files processed. Consequently, even if the user chooses search conditions that are time-consuming (e.g., many variable modifications, non specific cleavage, large FASTA file including isoforms etc.), the time price for that more complex search is paid only once, and not ‘n’ times for ‘n’ LC-MS files. Particularly for the larger projects (e.g., 100s or even 1000s of files), this is a significant time savings due to GH’s fundamental architecture.

We summarize all the above steps through a flow-chart (and include some additional components not central to this paper) in Fig. 3 below:

**Fig. 3:**
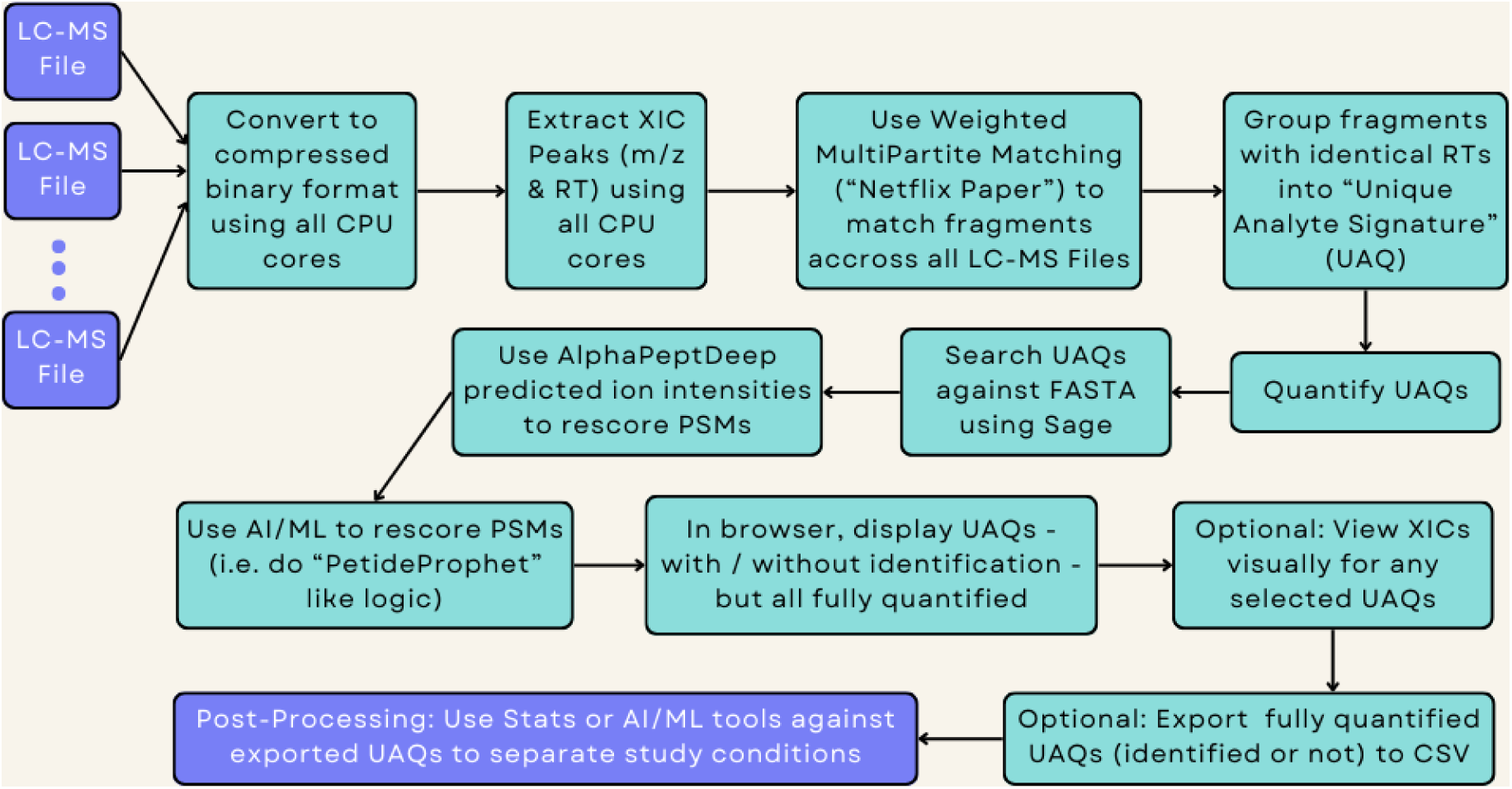
Flowchart Summarizing Core GH Steps. Purple boxes refer to input or post-processing steps outside of GH; green boxes refer to core GH steps.

## III. Non Computational Methods

One primary project, “AstralDilutionSeries55”^23^, and data subsets from three supporting projects (“AstralPlasma60”, “CovidPlasma61”, and “HeartTissue57”), were run on either the Astral MS (ThermoFisher)^24^ or Orbitrap Exploris 480 (ThermoFisher) and then analyzed by both GH and DIA-NN. The specific method details for the projects are described further below.

In the “AstralDilutionSeries55” project, the goal was to determine GH’s linear quantitation response for not only the ∼5-25% analytes identifiable in FASTA search spaces, but more importantly for the ∼75-95% of analytes in DIA-MS datasets (which contain the unexpected sequences or unexplored PTMs) that were not identifiable in FASTA search spaces. We used a Thermo Astral MS coupled to a Vanquish-Neo LC, over a gradient of 24 minutes, with 200 MS2 windows of 3 Th each, no overlap between windows, and DIA MS2 windows spanning the 380.4 to 980.7Th range (e.g., 380.4-3833.4Th, 383.4-386.4Th, etc.), as previously reported in in Fu *et al.*^23^ (MassIVE# MSV000094136). The MS1 was run at 240k resolution on the Orbitrap portion of the Astral, while the Astral Detector resolution for MS2 80k FWHM resolution at m/z of 524. We selected one pooled human plasma sample to run on the LC-MS 55 times at differing concentrations. Specifically, there were 11 groups of 5 technical replicates, and the 11 concentrations were 1000ng, 800ng, 600ng, 500ng, 400ng, 300ng, 200ng, 100ng, 50ng, 25ng, and 12.5ng. Search was performed against a human FASTA file with 1 missed cleavage allowed, and charges 1 to 5 inclusive considered. We used Sage v0.14.7^25^ and AlphaPeptDeep v1.0.1^26^ with GH; and for DIA-NN, we used v1.80. Although newer versions of DIA-NN are now available, at the time that the bulk of the work in this paper was performed, the production cluster had v1.80 installed. (We typically do not change the software on the production cluster quickly, and the number and sizes of the LC-MS files were such that attempting to run DIA-NN on a user’s desktop would not have produced results in any reasonable time frame.) That said, based on the readme documentation of the changes made in 1.8.1 or 1.9.0, we do not expect the central conclusions in this paper to be substantially different were version 1.8.1 or 1.9.0 to be used, particularly since we would explicitly not want to use “channel-q” logic (for SIL searches during the WOE calculations) for the reasons outlined in Note S3.

For the “AstralPlasma60” project, the LC, MS, and informatics search methods were identical to the “AstralDilutionSeries55” project, but there were 500 distinct patient plasma samples (as opposed to 1 pooled plasma sample at 11 differing loads), though for the purposes of this proof-of-principle paper, we used 60 of the 500 samples (roughly every 8^th^ run) for GH’s analysis (MassIVE# MSV000094136^23^).

For the “CovidPlasma61” project, there were 500 different human plasma samples run, but as with the previous project, we also used a subset (i.e., 61 samples) for GH’s analysis. These CovidPlasma61 samples were run on a Orbitrap Exploris 480 MS (ThermoFisher) coupled to a Evosep One LC on a 25 minute gradient, with MS1 resolution of 120k and MS2 resolution of 50k, and 50 MS2 DIA Windows of 22Th covering 349.5 to 1400.5Th, with 1Th overlap between windows (e.g,. 349.5 - 371.5Th, 370.5Th - 392.5Th, etc.). To compare results against standard DIA algorithms for identified-in-FASTA file peptides, we used a combination of MSFragger v3.4^27^ plus DIA-NN v1.80.

Finally, for the “HeartTissue57” project, we analyzed 57 patient heart biopsy tissue samples via GH and DIA-NN. The samples were run on a Thermo Orbitrap Fusion Lumos Tribrid coupled to a Dionex Ultimate 3000 nano LC on a 90 minute gradient, with MS1 resolution of 120k and MS2 resolution of 30k, and 150 MS2 DIA Windows of 4Th covering 400 to 1000Th, with no overlap between windows (e.g., 400-404Th, 404-408Th, etc.), as described further in Meddeb *et al.*^28^. The same setup of MSFragger 3.4 & DIA-NN 1.80 was used as with the CovidPlasma61 project.

## IV. Results & Discussion

For the primary project, “AstralDilutionSeries55”, we measured the R^2^ correlation values using GH’s algorithm’s *non-normalized* quantitation values and the expected concentration for the 55 samples (11 different concentrations * 5 technical replicates/concentration). In real biomarker projects, we would use the normalized quantitation results, as normalization reduces variance by ∼10%, but since this was a dilution series experiment and GH treated all 55 files as a single experiment, GH did not perform any normalization. (DIA-NN, however, was run in 11 independent batches of 5 technical replicates, and so normalization was performed within each batch.) We first did the correlation analysis for six peptides that (a) were preselected by the lab due to high biological interest and (b) had known high correlation R^2^ values in DIA-NN (Table 1).

**Table 1:** Linear R^2^ Values for 6 Preselected Peptides of Interest.

| Sequence | Charge | GH $R^2$ | DIA-NN $R^2$ |
| --- | --- | --- | --- |
| CEACPPGYSGPTHQGVGLAFAK | 3 | 0.98 | 0.99 |
| ESDTSYVSLK | 2 | 0.97 | 0.99 |
| IADVTSGLIGGEDGR | 2 | 0.97 | 0.99 |
| ILEGFQPSGR | 2 | 0.98 | 1.00 |
| SDVVYTDWK | 2 | 0.99 | 0.99 |
| YVGGQEHFAHLLILR | 2 | 1.00 | 1.00 |
| **Mean:** |  | ~0.982 | ~0.993 |

Correlation values for 6 peptides of interest to the lab for both DIA-NN and GH's algorithm, and which were previously selected by the lab to be of high quality from DIA-NN's perspective.

For these selected six peptides, we achieved a mean correlation R^2^ of 0.982, which was comparable to DIA-NN’s mean correlation R^2^ of 0.993, even though the quantitation calculations for GH’s algorithm were done without help from any useful information that we could gleam from the FASTA file, such as knowledge of the expected b or y ions, and DIA-NN, unlike GH, performed normalization within each of the 11 batches of 5 technical replicates. (If we were to add information from the FASTA file or do normalization within each batch, the quantitation could be improved even further; however, it’s extremely unlikely that improvements of ∼1% R^2^ for quantitation would make any major difference in *discovery*-proteomics studies. Moreover, these six peptides had been preselected as an example based on their high scoring R^2^ values when searched using DIA-NN.)

For those six peptides, we also visually inspected the relationship between GH’s algorithm’s non-normalized quantity vs. the actual concentration (Fig. 4A and Fig. S1-S5).

**Fig. 4:**
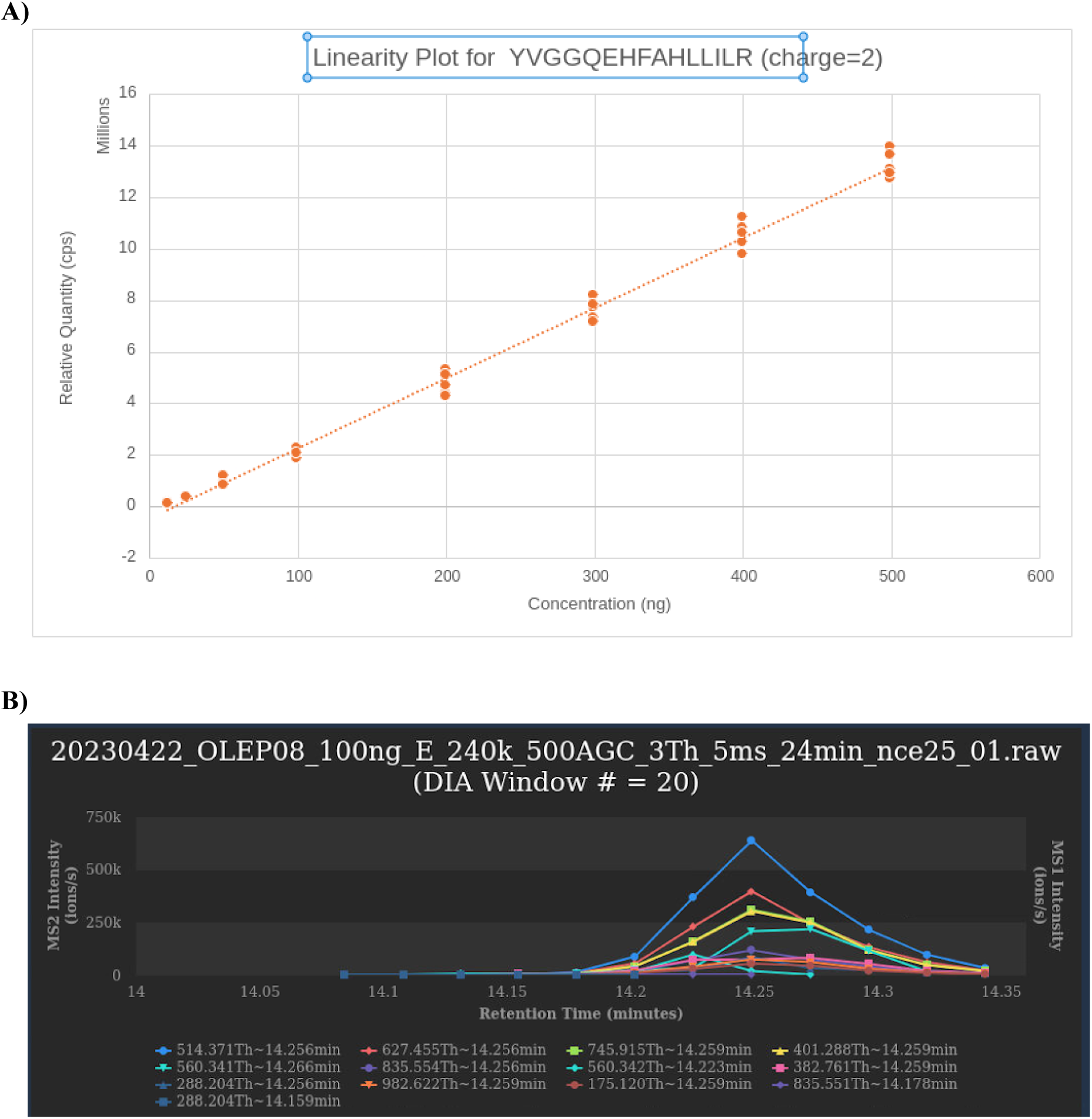
Linearity and XIC Plots for YVGGQEHFAHLLILR. **A)** Linearity plot for YVGGQEHFAHLLILR at charge state 2 for the range of concentrations where the relationship was expected to be linear (12.5ng to 500ng). **B)** The XIC plot for the YVGGQEHFAHLLILR peptide at the 500 ng concentration. Each colored curve represents an m/z fragment associated with this peptide, with the curve’s name defined as the m/z value (in Thomsons) followed by “∼” and then followed by the calculated interpolated retention time (using Savitzky-Golay smoothing) for the curve’s apex.

Based on the plots of all six peptides (Fig. 4A & S1-S5), we visually intuited that were a real biomarker project (e.g., hundreds of samples per study condition instead of the five samples per concentration) to have even only a ∼2x difference in abundance for a certain peptide (and assuming only modest variance), it should be relatively easy for both statistical routines (e.g., student t-test) or machine learning programs (e.g, XGBoost^29^) to pick out those peptides as peptides that differentiate one group of samples from another. Finally, we also plotted the XIC traces for these peptides and manually verified them. We show in Fig. 4B above the “YVGGQEHFAHLLILR” peptide’s XIC plots for one concentration, and we show similar plots for the two other concentrations in the supplemental (Fig. S6-S7). As expected, the MS2 XIC traces for the top expected fragments appeared in the most abundant samples (1000ng, Fig. S6), the medium-level abundance (100ng, Fig. 4B), and the least abundant concentrations (12.5ng, Fig. S7), all at roughly the same RT of 14.25 minutes, but with different MS2 intensities corresponding to the different concentrations.

Next, instead of focusing on only six peptides that were preselected from DIA-NN’s perspective as high R^2^ scoring peptides, we then calculated the percentage of *all* peptide spectrum matches (PSMs) identified at 1% false discovery rate (FDR) that had R^2^ values at or above 0.90 using either algorithm. Our initial goal was to achieve a percentage that was ∼80% of DIA-NN’s percentage. However, the percentage of identified PSMs quantified at or greater than 0.90 R^2^ was higher in GH’s algorithm (83%) than in DIA-NN (75%), even though GH did not (a) perform normalization within each batch as was done with DIA-NN and (b) use any useful information (e.g., knowledge of b/y ions) from the FASTA file (Table 2 below, and Note S6 for detailed calculations). (The absolute number of PSMs was claimed to be higher in DIA-NN; however, in other datasets described in later sections, we evaluate the claim of these additional PSMs.)

**Table 2:** Percentage of PSMs Quantified with R^2^ >= 0.90.

| $R^2$ Correlation Range | GH | | DIA-NN | |
| --- | --- | --- | --- | --- |
|  | # Quantified | % Quantified | # Quantified | % Quantified |
| 0.90 to 1.00 | 5292 | **83%** | 6,611 | 75% |
| -1 to 0.90 | 1089 | 17% | 2,245 | 25% |

The percentage of PSMs with $R^2$ correlation coefficients at or above 0.90 was higher for GH's algorithm (83%) vs DIA-NN (75%), even though GH's goal was not to focus on identifiable peptides or to use information from FASTA files or spectral libraries to help with quantitation. (While the absolute number of PSMs was claimed to be higher in DIA-NN, we manually analyze in other projects the claimed additional DIA-NN PSMs with the lowest DIA-NN q-values.)

Finally, we focused our analysis on the key goal of GH: quantifying those peptides present in the MS but not necessarily identifiable in the FASTA-file search spaces. While 5,938 PSMs were identified at 1% FDR, 117,560 analytes (i.e., unidentified peptides at unknown charge states, plus an expected relatively small number of non-peptide artifacts, such as chemicals present in the MS room’s air etc.) were quantified-but-not-identified by GH’s algorithm, i.e., ∼95% of analytes in the MS were now quantifiable (117,560 / (117,560 + 5,938)) even though they were not found in the FASTA search space.

Since we pre-grouped the quantified analytes into three bins (“high-quality”, “medium-quality”, and “low-quality”) based on internal factors, we analyzed in Table 3 below the number of unidentified analytes that met or exceeded the same R^2^ correlation of 0.90 for each of those three quality bins:

**Table 3:** Number of *Unidentified* Analytes that were Quantified at R^2^ >= 0.90.

| *Unidentified Analytes Only* |  |  |  |  |  |  |
| --- | --- | --- | --- | --- | --- | --- |
|  | High-Quality |  | Medium-Quality |  | Low-Quality |  |
| $R^2$<br>Correlation | #<br>Quantified | %<br>Quantified | #<br>Quantified | %<br>Quantified | #<br>Quantified | %<br>Quantified |
| 0.90 to 1.00 | 33890 | **80%** | 27708 | 48% | 4014 | 23% |
| -1 to 0.90 | 8285 | 20% | 29771 | 52% | 13471 | 77% |

As expected, the high-quality analytes had the highest “% Quantified” for unidentified analytes of 80%, which was similar to the “% Quantified” of 83% for identified PSMs from the previous Table 2. Even though the medium and low-quality analytes had a lower percentage of analytes that met the expected quantitation R^2^ linearity analysis, we do not automatically discard those medium and low-quality analytes at this stage. Instead, we allow any downstream external statistical tool (e.g., student’s t-test) or machine learning / AI program (e.g., XGBoost or possibly neural networks) to automatically sift through the data so that it could separate the signal from the noise rather than prematurely discarding possibly valuable data.

One challenge with the “Jitter Deconvolution” algorithm described above is that it could be computationally demanding, since we attempt to match MS2 fragments between samples without relying on a prebuilt and comparatively small peptide library. And in general, for at least ∼5 years, if not the last 10+, the challenge of almost all these discovery-proteomics MS projects was the extensive computational time to process the large MS datasets, almost always exceeding the MS time. And, as MS instruments have evolved, they now produce file sizes that are ∼5x to ∼10x larger than the previous generation. Given the complexity involved in global deconvolution without use of a FASTA file or spectral library, our initial hope was that GH would take not too much longer than MSFrager+DIA-NN (e.g., 2x to ∼4x *slower* than MSFragger+DIA-NN). To our surprise, however, we were >40x (“theoretical MSFragger+DIA-NN speeds”) to >200x (“MSFragger+DIA-NN speeds under real load conditions”) faster than MSFragger+DIA-NN (Fig. 5); and compared to the LC-MS runtimes, we were ∼10x faster than the Orbitrap Exploris 480 MS (i.e., for the CovidPlasma61 project, it took 2.2hrs for GH’s algorithm vs 24.5 hours for the Exploris MS) and comparable to the Astral (i.e., 23.8 hours for GH’s algorithm vs ∼24 hours for the Astral MS). (We define “load conditions” in Fig. 5 below.)

**Fig. 5:**
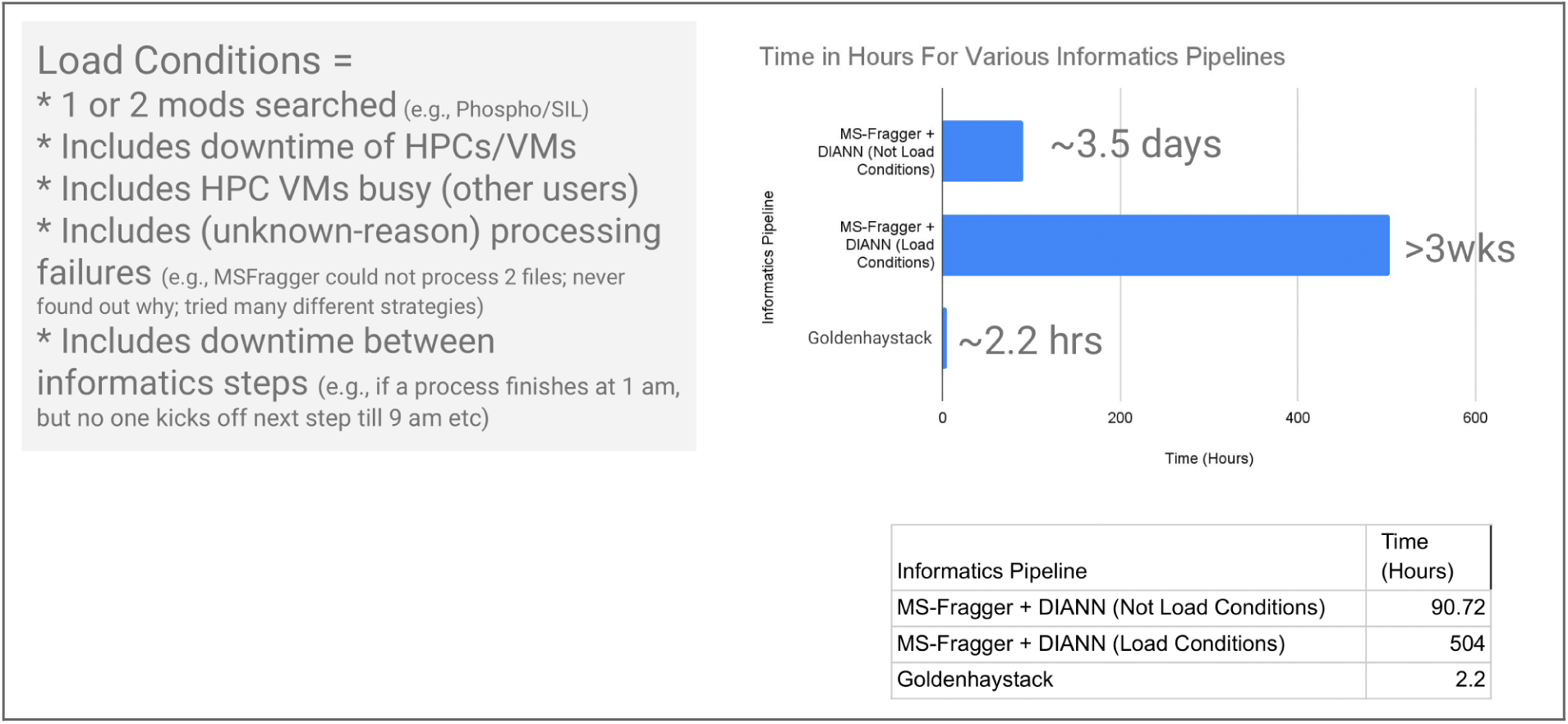
End-to-end Informatics Time Under Load Conditions. Runtimes for DIA-NN and GH’s algorithm as observed for the “CovidPlasma61” project. (In particular, during the informatics portion of the CovidPlasma61 project, real-life situations arose, such as high performance computing (HPC) system having unscheduled downtimes, or HPCs being busy due to use from other labs (e.g., genomics labs), or two MS files failing to process in MSFragger for no reason that we could resolve in any reasonable time etc.) We refer collectively to these scenarios as real life “load conditions”. GH’s algorithm ran in ∼2.2 hours end-to-end, which was >40x faster than MSFragger+DIANN’s theoretical top speeds (i.e., ignoring most load conditions, but including the variable PTM search parameters), >>100x faster than the real, observed runtime (which includes the load conditions), and >10x faster than the MS acquisition time.

GH’s speeds were achieved on a single, albeit very large, computer of 180 CPUs (360 virtual threads) and 1.4TB RAM. So, if needed, a cloud-computing version of GH, which could use a nearly unlimited number of computers in the cloud, could be developed to process the datasets even faster.

The above sections describe ∼85% of the core of GH’s motivation, algorithm, and primary benefits, and so one can safely skip to the conclusion. That said, we also ran GH’s algorithm against three other DIA-MS datasets, all of which were different along at least one key dimension, e.g., differing biofluids, tissue types, LC types, LC gradient lengths, or MS instrument types, even though there were no theoretical reasons to suggest that the “Jitter Deconvolution” technique would not work under these different instrument or sample conditions. Each project typically provided one interesting insight, as described further below.

For the “CovidPlasma61” project, which used the Orbitrap Exploris MS and 61 plasma samples from different humans (instead of the same sample run 5 times at 11 different dilutions), we discovered that we not only identified more synthetic, spiked-in iRT^30^ peptides (11 distinct peptides out of a total of 11 spiked in, whereas DIA-NN found only 10 out of 11 peptides) (Table 4 below), but it also correctly claimed far fewer never-spiked-in stable isotopically labeled^31^ (SIL) peptides (i.e., peptides with a heavy label on K or R) compared to DIA-NN, which claimed 237 SIL peptides, a 11750% increase over GH’s claim of 2 non-existent SIL matches. The correct answer for the number of SIL peptides is 0, but acceptable answers could be between 0 peptides and the number of peptides corresponding to a ∼1% FDR rate.

**Table 4:**
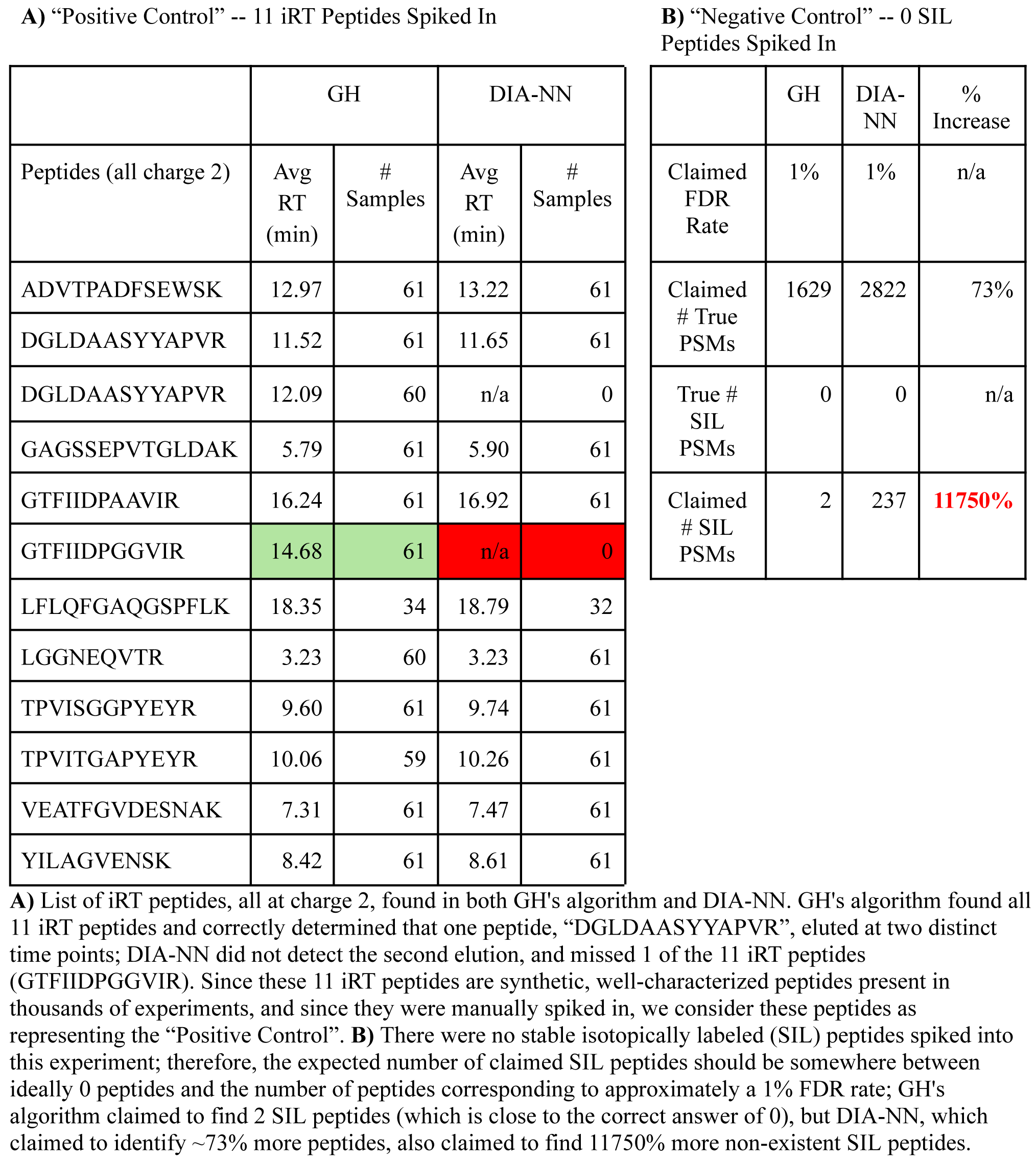
iRT & SIL Peptides Analysis.

If we think of the analysis of iRT peptides as “positive control” -- since we know that we spiked in 11 iRT peptides, and these 11 iRT peptides are controlled, synthetic peptides that have been spiked into thousands of projects in other labs -- and we think of the SIL peptides as “negative control” -- since there were 0 SIL peptides spiked in, and the SIL modification does not occur in nature -- then even for these “identified purely through FASTA files” peptide analysis, GH’s algorithm showed better performance for the positive control and much better performance for the negative control. This was a surprising result, as the focus of GH’s algorithm was not identifying peptides from FASTA files, but instead it was to quantify the up-to 95% of peptides present in the MS that are not identifiable in the FASTA file. Further, although we were able to identify and quantify 2,388 PSMs, we quantified a total of 33,138 unidentified analytes, i.e., ∼93% of analytes in the DIA-MS dataset (33,138 / (33,138 + 2,388)) were now quantifiable even though they were not found in the FASTA search space for this “CovidPlasma61” project.

For the “HeartTissue57” project, the main difference with the “CovidPlasma61” project was that it used tissue instead of plasma and the gradient length was 90 minutes instead of 25. While both DIA-NN and GH’s algorithm identified the expected iRT peptides and GH’s algorithm had a reasonable number of SIL peptides identified (close to the 1% FDR), DIA-NN claimed to identify 26,046 PSMs at 1% FDR vs. GH’s algorithm’s 13,271 identified PSMs. As we wished to understand why DIA-NN claimed to identify so many PSMs that GH claimed were not identifiable at each algorithm’s stated 1% FDR rate, we selected the PSMs claimed to be found in DIA-NN but not in GH’s algorithm, sorted them by DIA-NN’s q-value in ascending order (lower q-values means that DIA-NN is more certain it is a true match) and then, because many PSMs had identical low DIA-NN q-values of 5.28E-5, we sorted again by DIA-NN’s abundance metric in descending order, and then selected those final best 5 peptides (Table 5).

**Table 5:**
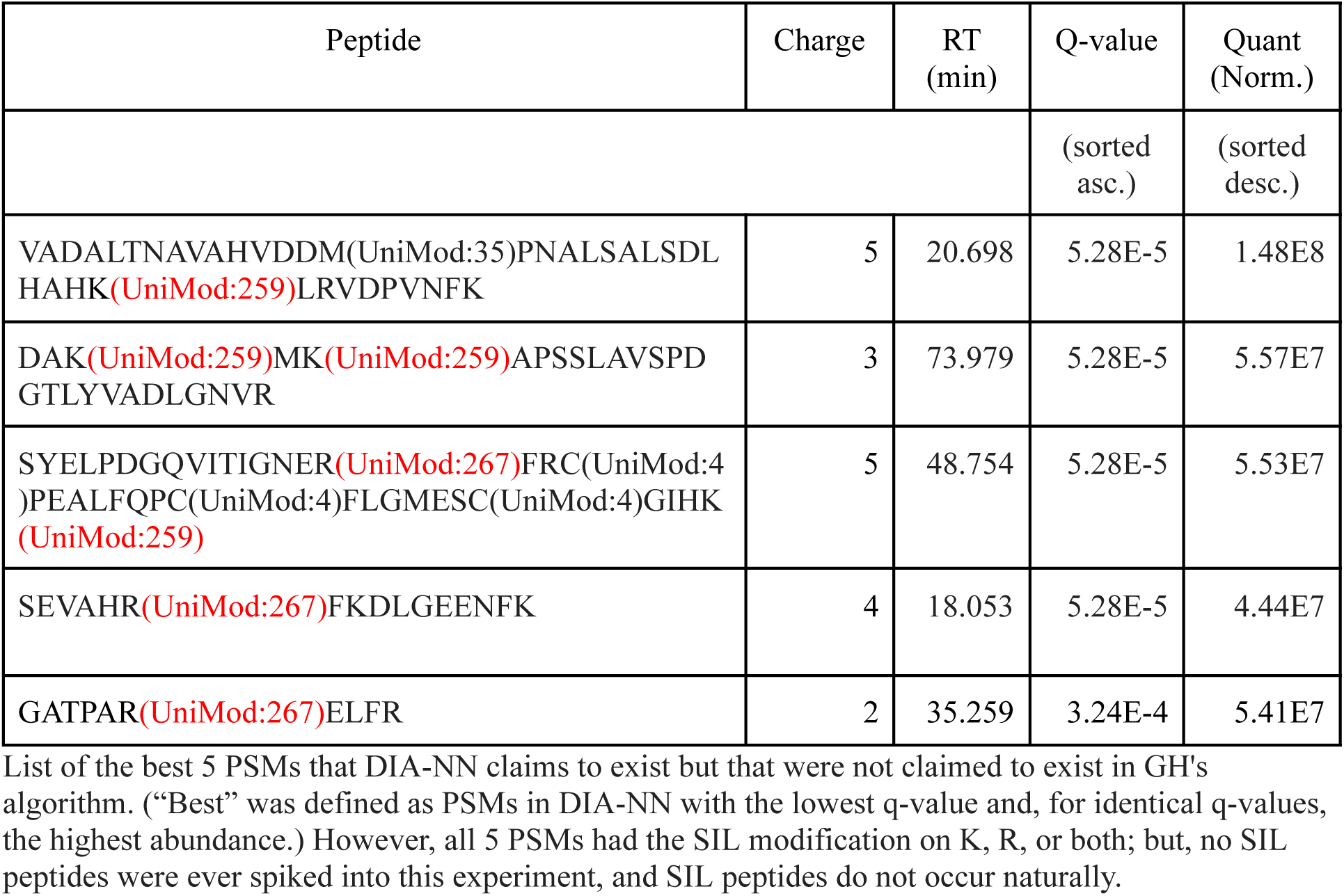
Top 5 PSMs in DIA-NN but not in GH’s Algorithm (HeartTissue57 Project)

Of these resulting best 5 PSMs claimed to exist by DIA-NN but not by GH, all contained a SIL modification on either K, R, or sometimes both (red text in Table 5). Since no SIL peptides were spiked in, and since SIL modifications do not occur naturally, these “best 5” PSMs found in DIA-NN but not GH’s algorithm were false matches. Finally, GH’s algorithm quantified 40,208 analytes that were not identified in the FASTA search space, i.e., ∼75% of analytes (40,208 / (40,208 + 13,271)) in the MS were now quantifiable even though they were not found in the FASTA search space for the “HeartTissue57” project.

Finally, the last project we considered was the “AstralPlasma60”. This dataset was similar to the first one, “AstralDilutionSeries55”, except that there were 60 different human plasma samples, instead of a single pooled sample run 5 times for each of 11 different concentrations. Moreover, we searched this project with the same variable modifications as with CovidPlasma61 and HeartTissue57, i.e., oxidation on M, phosphorylation on S, T, or Y, and SIL modifications on K or R, even though no SIL peptides were spiked in. The key goal was to ensure that the algorithm simply ran and returned reasonable results, as the total size of the Astral datasets, even for merely 60 samples, were humongous (> 350GB). From beginning-to-end, it took 23.4 hours to process all 60 files compared to ∼24 hours for the LC-MS, and GH’s algorithm found all 11 iRT peptides in all 60 samples, claimed to find 95 SIL peptides (which is close to ∼1% FDR), found 6,495 peptides overall at a 1% FDR rate, and quantified 121,969 analytes that were not identified. Unfortunately, while DIA-NN could be run without phosphorylation and SIL modifications, it could not be run in any reasonable timeframe with those modifications included, even on a cluster, and so could not be used to compare the results with GH’s algorithm as was done for the CovidPlasma61 and HeartTissue57 projects which had much smaller LC-MS file sizes. The net conclusion we drew from this project was that (a) Astral is indisputably an excellent MS with noticeable improvements in final peptide results compared to Exploris, (b) but it does so at the cost of extraordinarily large datasets (e.g., ∼8GB / file, and this too just for the less complex plasma samples) and (c) and despite the large data sizes per run and our desire to still be able to search for variable PTMs, we could still process those large datasets slightly faster than the MS with the current architecture (i.e., without customizing GH’s algorithm to run on hundreds of computers on the cloud) while quantifying the ∼95% of the analytes in the Astral MS (121,969 / (121,969 + 6,495)) that were not found in the FASTA search space.

## V. Miscellaneous

Although throughout this paper we have specified 50 files as the minimum number of files for GH to apply the “Jitter Deconvolution” algorithm (and all four datasets in this paper used roughly that number of MS files in each project), the “Jitter Deconvolution” may still work reasonably well with fewer number of LC-MS files, especially on datasets in which the samples belong to different individuals rather than the same sample analyzed multiple times but with slightly different concentrations, as is common in mixed matrix samples. (Real samples from different individuals are more likely to have LC jitter as the samples contain different peptides in different concentrations, so there will be more likelihood of matrix effects and hence LC jitter that GH can take advantage of.) In other words, the “Jitter Deconvolution” tries to *gracefully* degrade as the number of samples ‘n’ tends towards a hard lower limit of 15. Conversely, “Jitter Deconvolution” modestly improves as ‘n’ exceeds 50 (e.g., ∼5-10% improvement in quantifiable analytes when ‘n’ is somewhere between ∼100 and ∼200), but starts to have limited additional benefit (e.g., <1% improvement in quantifiable analytes) as ‘n’ >> ∼200. That said, although outside the needs of GH’s “Jitter Deconvolution” logic, we suspect that external statistical and especially AI / ML tools (which are post-processing tools that can be used to determine a panel of peptides that separates study conditions) would see substantial benefit for many projects when ‘n’ is well beyond ∼200 or even ∼1000. That is, for some projects, one may be lucky and not need a large ‘n’, e.g., the biomarker to determine pregnancy may be a single simple protein with a nearly clear step-wise, binary response, and so ‘n can be small; conversely, other studies may have a less binary, more subtle and on-a-continuum-like response based on a multitude of peptides, and so larger ‘n’ would be needed to achieve a more subtle separation.

## VI. Limitations

While GH can quantify both the ∼5-25% of peptides in DIA-MS datasets that are identifiable in the FASTA search space as well as the ∼75-95% that are not in the FASTA search space, GH does not address the “dynamic range”^32,33^ problem. That is, the dynamic range between the most abundant peptide and the least abundant peptide is high (∼6 to ∼7 orders of magnitude for cells^32^ and in excess of ∼10 orders of magnitude for plasma^34^), whereas the best LC-MS’s discovery proteomics detection abilities have a dynamic range of ∼4 to ∼5 orders of magnitude^24^. For those projects in which there is a high likelihood (e.g., > 50%) that a parsimonious peptide panel that separate study conditions exists within the LC-MS’s dynamic range, we should be able to find that peptide panel soon, though validation would still take months to years; for projects in which the likelihood of a peptide panel being within the LC-MS dynamic range is low (e.g., ∼1 to ∼5%), it would take >>10 projects to ensure a reasonable likelihood that one project will succeed in providing us with a peptide panel, even though the success of a single project may easily outweigh the efforts of dozens of other “failures”; however, for those projects in which the likelihood of a peptide panel being within the LC-MS dynamic range is very low (e.g., <<1%), we may need orthogonal techniques to *circumvent* the dynamic range problem altogether (e.g., Mag-Net^35^, Seer^36^, PTM enrichments, PreOmics^37^, and the recently published perchlorate acid (perCA) method^38^ etc.) Further, we may also give slightly more weight to studies in which there is a higher likelihood that a peptide panel is within the LC-MS’s dynamic range for the tissue type being examined (e.g., almost any disease involving tissue cells, e.g., skin cell biopsies for skin carcinoma, lung cell biopsies for non-small cell lung carcinoma etc., or, for plasma, the approximate top third of diseases described in Fig. 4c in a recent plasma proteomics MRM/Olink paper^39^). Irrespective of which of the above scenarios we find ourselves in, we look forward to the next set of computational challenges.

## VII. Concluding Remarks

Using four different datasets from two different instruments, we tested the following idea: GH can separate MS2 fragments into their unique analyte signatures and then quantify them accurately through the “Jitter Deconvolution” strategy. That is, since peptides that exactly co-elute in one subset of samples do not exactly co-elute in another subset of samples (the “jitter” in LCs), GH can (a) match MS2 fragments across all samples using a weighted multi-partite matching algorithm (the “Netflix” paper), then (b) computationally separate MS2 fragment ions, then (c) regroup the MS2 fragment ions into unique analyte signatures while simultaneously reducing stochastic noise (since stochastic noise is random and will not be consistent across a sufficient number of samples), then (d) quantify those unique analyte signatures (without being limited to quantifying only the ∼5-25% of peptides in DIA-MS datasets that are in a FASTA or spectral library search space), and then (e) finally identify those spectra that can be matched to a predicted spectra in the user-supplied FASTA search space.

Consequently, GH not only quantifies and identifies the ∼5-25% of unique analyte signatures in DIA-MS datasets that are identifiable in the FASTA search space – and does so with as much as ∼11270% fewer false SIL matches and comparable or occasionally slightly higher number of iRT identifications than the leading FASTA-only DIA-MS algorithm (MsFragger + DIA-NN) – but more importantly GH quantifies the remaining ∼75-95% of unique analyte signatures present in DIA-MS datasets that were previously not quantified. Further, from an operational perspective, GH can process DIA-MS datasets at ∼1x (for Astral data sets) to ∼10x (for non Astral Thermo data sets) faster than the LC-MS acquisition time, and for non-Astral datasets, between ∼40x to ∼200x faster than the leading FASTA-only DIA-MS algorithm. (For Astral datasets, MSFragger+DIA-NN could not process the files within any reasonable timeframe when considering three variable modifications.) We summarize in Table 6 the quantification, quality, and runtime performance metrics for all four DIA-MS datasets.

**Table 6:** Summary of Quantification, Quality, and Runtime Performance Metrics.

| Data Set<sup>(a)</sup><br># | GH Quantification Metrics: |  |  | Quality Metrics: |  |  |  | Runtimes (In Hours): |  |  |
| --- | --- | --- | --- | --- | --- | --- | --- | --- | --- | --- |
|  |  |  |  | GH |  | DIA-NN<sup>(b)</sup> |  |  |  |  |
|  | # PSMs In FASTA | # Analytes Not In FASTA | % Analytes Not In FASTA | # iRT | # SIL | # iRT | # SIL | LC-MS | GH (on a single 360 core computer) | DIA-NN (on cluster) no load conditions<sup>(c)</sup> |
| 1 | 5,938 | 117,560 | 95 | 11 | n/a<sup>(d)</sup> | 11 | n/a<sup>(d)</sup> | ~24 | 22.8 | ~68 |
| 2 | 2,388 | 33,138 | 93 | 12<sup>(e)</sup> | 2 | 10<sup>(f)</sup> | 237<sup>(g)</sup> | ~24 | 2.5 | >90 |
| 3 | 13,271 | 40,208 | 75 | 11 | 204 | 11 | 653 | ~86 | 4.1 | >90 |
| 4 | 6,495 | 121,969 | 95 | 11 | 95 | (could not run)<sup>(h)</sup> |  | ~24 | 23.4 | (could not run)<sup>(h)</sup> |

In terms of possible future research direction, the “Jitter Deconvolution” algorithm could be generalized for other analytes that can be ionized in the LC-MS, as it does not need or even use the FASTA or spectral library search spaces at all. That is, GH could be applied to not just peptides / immunopeptides / glycopeptides (the proteomics family of analytes), but also to lipids, metabolites, cannabinoids, drugs-of-abuse, food additives, pharmaceutical drugs, and many other LC-MS ionizable analytes for which there is value in knowing what parsimonious set of analytes – initially identifiable or not – separates two or more study conditions.

Finally, as the ∼75-95% set of quantified-but-not-found-in-FASTA-search-space peptides contain, by definition, the *unexplored* PTMs (e.g., glycosylation, citrullination, ubiquitination, acetylation, phosphorylation, methylation, sumoylation, and hundreds more) and *unpredicted* peptide sequences (e.g., splice variants, sequence variation, protein precursors, unusual proteolytic cleavages, novel proteins, etc.), we suspect that this ∼75-95% set has a higher likelihood of differentiating study conditions (ref. ^40,41^ and Note S5) compared to peptides found in (a) typically generic (i.e., not typically disease or cohort specific), (b) genomics-derived (i.e., known-to-be-incomplete) FASTA search spaces with (c) only one or two PTMs considered (i.e., instead of hundreds). And, we only need one, two, or a small few of these unpredicted peptide sequences or unexplored PTMs to potentially create a parsimonious peptide panel that separates study conditions and that therefore possibly makes a real-world difference^42^.

## Data Availability

All DIA-NN and GH results for all four projects, plus the Thermo MS raw files for the “CovidPlasma61” and “RawHeartTissue57” projects, have been uploaded to MassIVE under the identifier MSV000096666 and ftp access of ftp://massive.ucsd.edu/v07/MSV000096666/; for the AstralDilutionSeries55 and AstralPlasma61 projects, the Thermo MS raw files can be found in MassIVE under identifier MSV000094136 and MSV000094137 respectively.

## Author Contributions

GS designed the algorithm and wrote the paper; QF and AB helped edit the paper and ran the mass spectrometers and the DIA-NN software program (QF worked on “AstralDilutionSeries55” and “AstralPlasma60” studies; AB worked on “CovidPlasma61” and “HeartTissue57” studies); JVE managed this project and edited the paper.

## Acknowledgements

We would like to thank Vadim Demichev, developer of DIA-NN, for helpful and informative discussions regarding how DIA-NN calculates q-values for peptides (irrespective of whether they have variable modifications).

