## Supplemental 1 for "Quantifying the ∼75-95% of Peptides in DIA-MS Datasets that were not Previously Quantified"

This document is divided into three sections. The first section includes all the supplementary figures (Fig S1 - S7); the second section includes all the supplementary notes (S1 - S6); and the third and last section includes citations for references that appear in this supplemental.

**<u>Supplementary Figures:</u>**

**Fig. S1: Linearity Plot for ESDTSYVSLK**

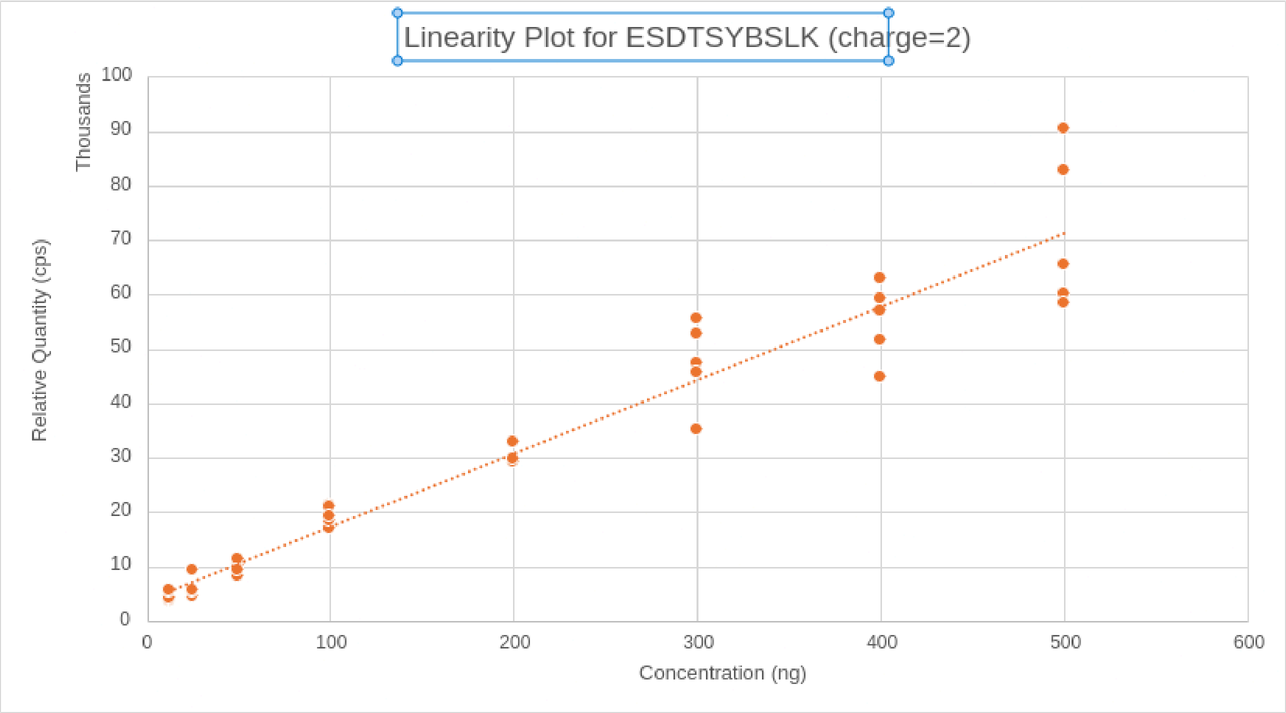

**Fig. S2: Linearity Plot for IADVTSGLIGGEDGR**

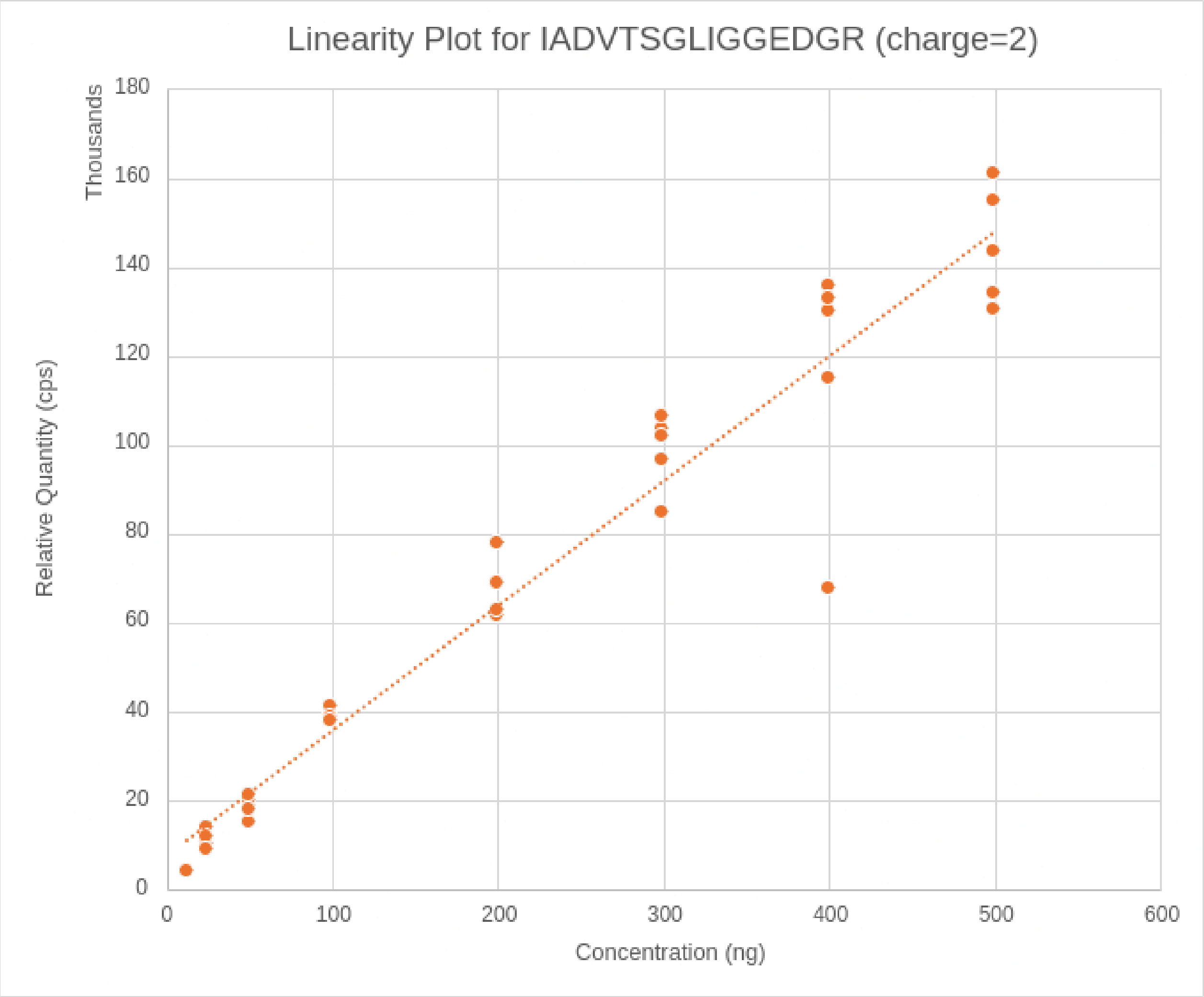

**Fig. S3: Linearity Plot for ILEGFQPSGR**

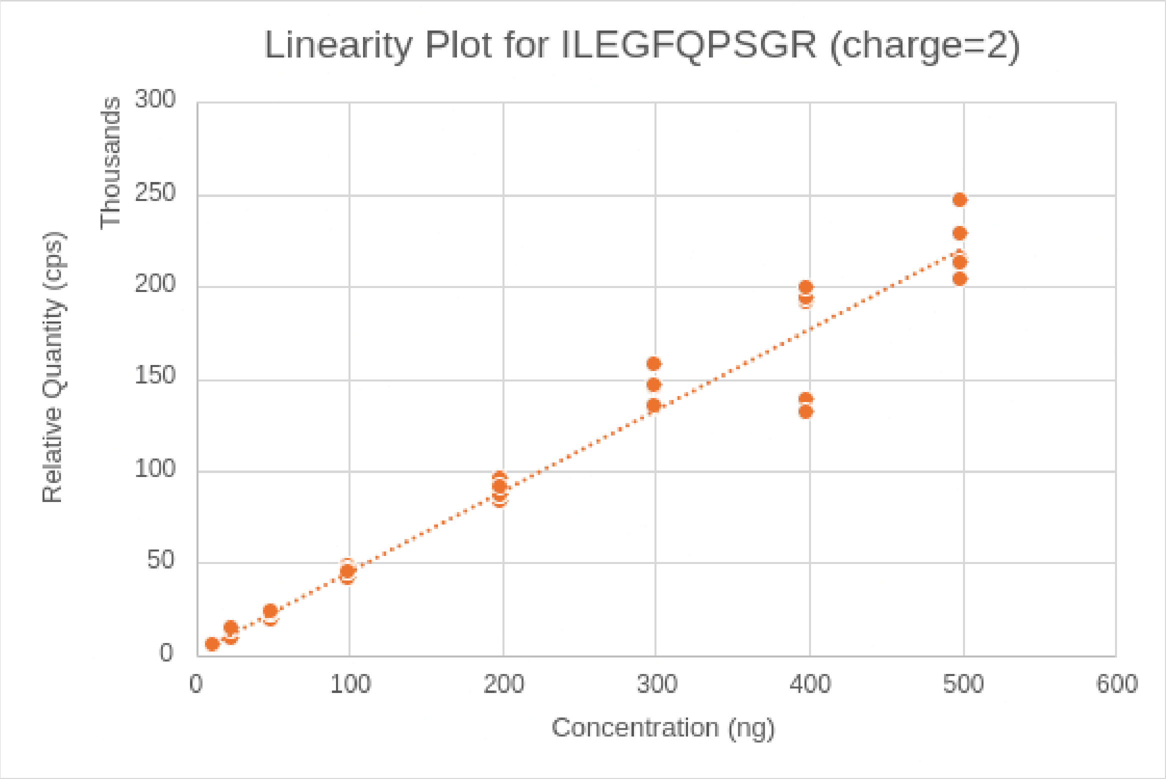

**Fig. S4: Linearity Plot for SDVVYTDWK**

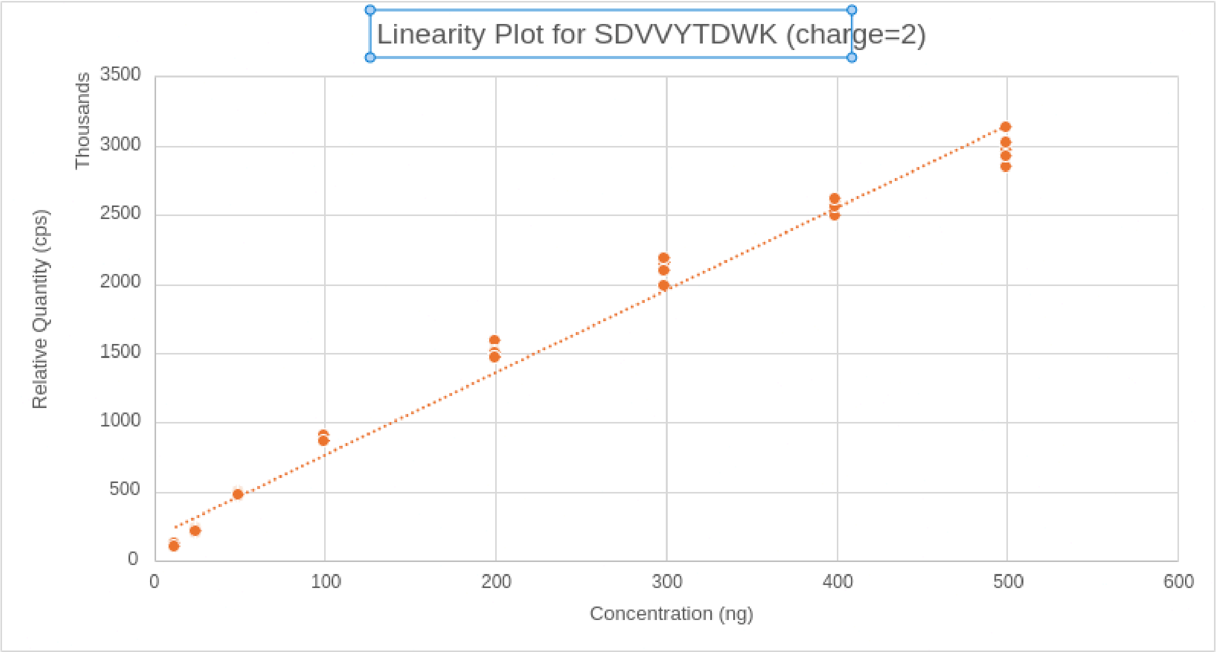


**Fig. S5: Linearity Plot for CEACPPGYSGPTHQGVGLAFAK**

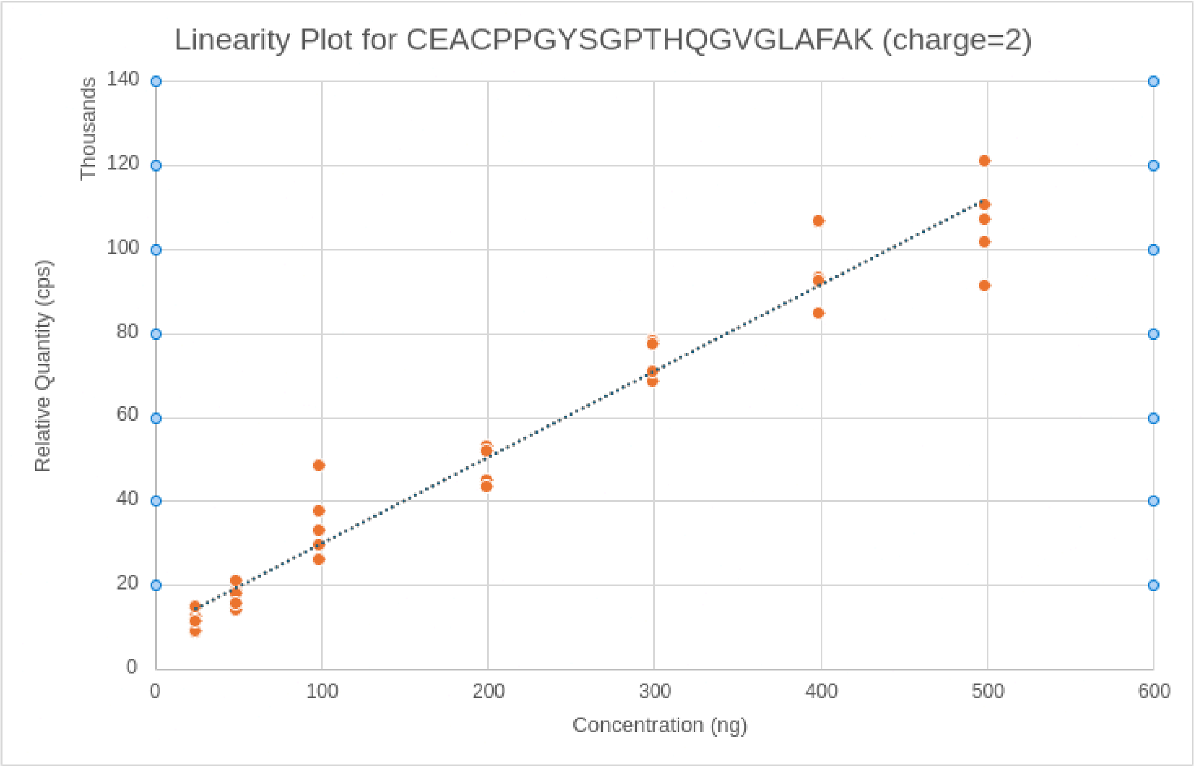

**Fig. S6: XIC Plot for YVGGQEHFAHLLILR at 1000ng concentration**

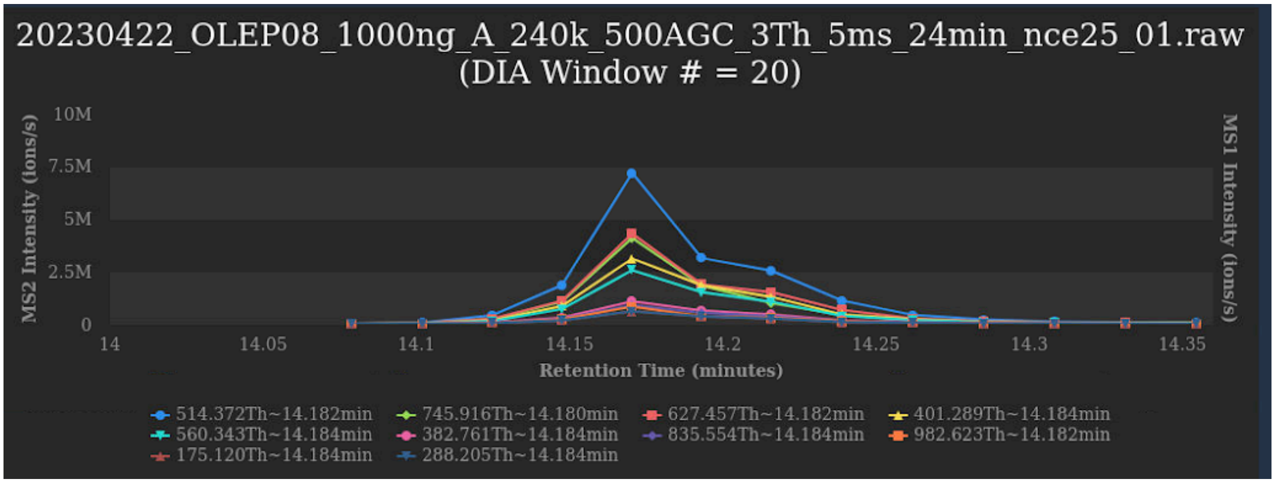


**Fig. S7: XIC Plot for YVGGQEHFAHLLILR at 12.5ng concentration**

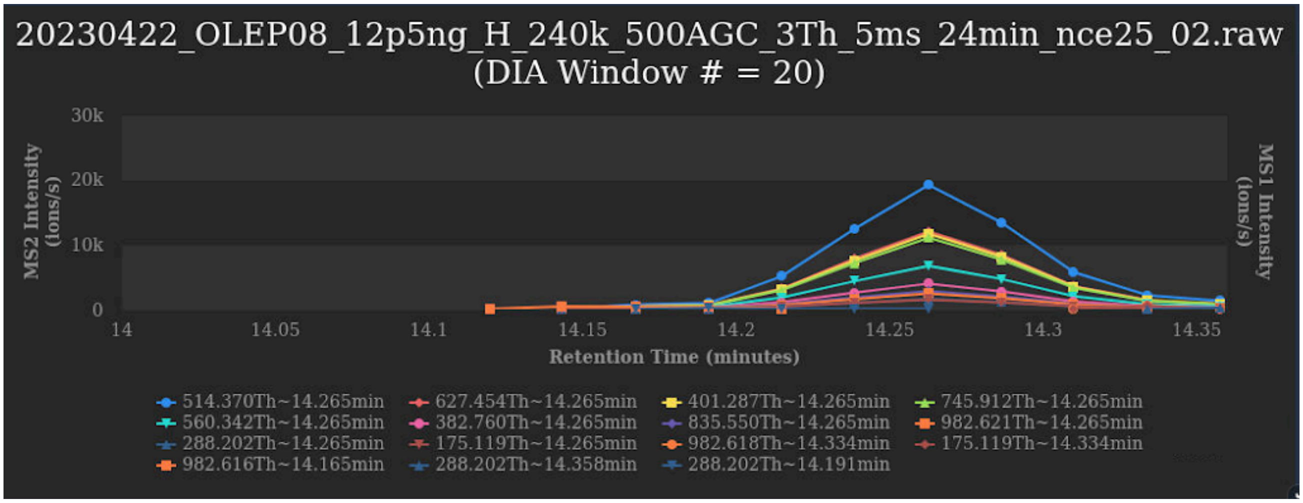


<u>**Supplementary Notes:**</u>

**Note S1:**

Even with DDA, where data is considerably less chimeric than DIA, denovo sequencing has not gained nearly as much traction as FASTA-based searching algorithms. User's reluctance in using de-novo algorithms for either DDA or DIA data may be due to the very long time it typically takes for de novo sequencing DIA algorithms to process DIA datasets, even substantially exceeding the FASTA-based algorithms such as DIA-NN, which themselves are taking too long (please see Fig. 4 in main paper). Moreover, the pre-built de novo sequencing algorithms typically only support less than half-a-dozen variable modifications, and training a new de novo model for newer PTMs would take a tremendous amount of time, a reasonable amount of bioinformatics expertise, and would require a substantial number of those PTMs to be first identifiable in the training datasets to begin with in order to ensure quality training. Further, the quality of denovo, owing to its much more complex problem that it is trying to solve, is not as high as FASTA-based algorithms. That is, it misses a substantial number of peptides that FASTA-based algorithms discover. Finally, the number of additional peptides that de novo identifies is modest (eg ~2x the number of peptides identified by FASTA-only algorithms), whereas the number of quantifiable analytes present in the DIA-MS datasets is up to 20x more, so the gain achieved by de novo algorithms may not be sufficient to overcome the aforementioned user concerns.

That said, we may, in future research, use DDA denovo algorithms, such as Cassanova[1], in a different manner, as follows: once all peptides in a DIA-MS datasets have been quantified by GH and then analyzed through some post-processing statistical (e.g., R) or AI (e.g., XGboost) program and users then therefore have a *small (e.g, << 10)*, parsimonious set of peptides – the "peptide panel" – that separates study conditions, it could be very useful to have a DDA de novo algorithms such as Cassnova provide a list of "top n" (where n could be high, e.g., 100) possible peptide sequences generated for those few peptides comprising the "peptide panel". Since Cassonava would be applied only on a very small number of peptides (e.g, << 10) which comprised the peptide panel instead of tens of thousands of possible peptides in the entire DIA-MS dataset, the de novo programs could take as much time as it wishes to do sophisticated analysis for those limited ~10 peptides of interest. Then, a mass spectrometrist trained in the classical ways of interpreting spectra could use the Cassonava provided list of possible sequences for any given "unknown" spectra as a starting point to identify / confirm an unknown analyte's proposed amino acid plus possible PTM sequence.

**Note S2:**

There are a number of differences between the "overlapping window" approach by the MacCoss lab and the "n-Offset Windows" approach described in the main paper. (As a side note, as mentioned in the main mapper, we ultimately do not use the n-Sample Offset approach in GH.) First and foremost, although in theory the "overlapping window" approach can have three overlapping windows, in practice, it is limited to only two overlapping windows, which means that at most the calculated effective window size can be reduced by 2, e.g., the effective computed window width is, at best, ½ the original window width. In contrast, the "n-Sample Offset" window approach can have "n" different starting-stopping windows (where "n" corresponds to the number of MS files in an experiment), which allows a reduction of the effective computed window size to 1/n of the original window size. (In practice, however, for other physical reasons related to the sharpness of the quadrapole, the effective window size is limited to being no less than ~0.07Th.) The second difference is that the "overlapping window" approach is, to the best of our knowledge, patented and the patent is currently owned by Thermo, and so is not, to the best of our knowledge, available on any other MS platform. In contrast, the "n-Offset Windows" is a technique that does not depend on any changes to the MS (in fact, the MS is running, as far as it is concerned, with the exact same parameters / idealized specifications as before, except that the windowing is slightly different from sample to sample). Consequently, the "n-Offset Windows" approach works on any MS capable of the standard DIA-MS protocol. Finally, the "overlapping window" approach makes an implicit assumption, because of its use of non-negative least squares optimization, that the intensity in any two adjacent points in the RT dimension in an XIC is the same. This assumption is technically false, though it is a reasonable approximation in some cases. However, some XICs may be changing much faster (i.e., a narrow elution peak) than others, and so this assumption introduces errors that may ultimately impact quantitation accuracy. In contrast, "n-Offset Windows" makes no such assumptions and therefore does not suffer from the same approximation-related errors.

**Note S3:**

When performing "Within Organism Entrapment" (WOE), one must not supply any helpful information to any informatics algorithm regarding the extra, not-naturally-occurring variable modification. The only three facts that the informatics algorithm should be aware of are: (a) whether the modification is variable, (b) the specific amino acid(s) the modification applies to; and, (c) the mass shift of the modification. In particular, the informatics algorithms must not be aware that the WOE related variable modification is a SIL modification. To the best of our knowledge, the main DIA informatics algorithms do not have any built-in special detection and then handling of SIL modifications unless users explicitly enable "SIL" parameters for SIL-related logic, but doing so would defeat the purpose of the WOE test.

If users are concerned that future versions of existing informatics algorithms or wholly new future algorithms attempt to use knowledge of the SIL modification to somehow artificially reduce the number of SIL identifications (and thus break the point of the WOE test), users could perhaps choose a new mass shift or new amino acid for the WOE's variable modification.

Further, a more advanced version of the WOE test would include physically spiking in a small number of SIL peptides (e.g., 100-200) in the samples at reasonable concentrations (i.e., substantially above the LOQ) and then counting the number of correctly identified SIL peptides vs incorrectly number of claimed SIL peptides. This is a useful test in-and-of-itself, but it also helps minimize gaming the WOE test by future algorithms.

**Note S4:**

For traditional entrapment (TE), a bioinformaticist would first need to identify a different organism's FASTA (d-FASTA) file which has minimal sequence overlap with the normal FASTA (n-FASTA) file. Then, once a suitable organism FASTA file has been identified, the bioinformaticist would need to concatenate the d-FASTA with the n-FASTA file. Ideally, the bioinformatics would also eliminate any overlapping peptide sequences (while considering the missed cleavages) between the d-FASTA and n-FASTA files. After the search has completed, to correctly calculate the re-estimated FDR, the bioinformaticist would need to statistically correct for the difference in sizes between the n-FASTA and d-FASTA files. None of the above is overly complicated, but it still takes at least a few hours and involves multiple steps that may not be as trivial as they first appear (and therefore leaves room for error / variation between different bioinformaticists etc.)

**Note S5:**

Almost by definition, when running discovery proteomics studies, we are interested in what is particularly *distinct*, from a proteomics perspective, about a subset of people (e.g., obese, ~90-year old males who are infected with covid but display no or very mild symptoms vs their matched counterparts who respond poorly to covid infection; or, for example, ~25-year old, previously healthy female runners who are on ventilators following covid infection vs their matched counterparts who display no visible symptoms etc.) rather than the generic information (i.e., typically not cohort specific; typically not disease specific; and incomplete in general, due to the known imperfections in attempting to predict protein sequence from genomics data etc.) contained in FASTA files. Consequently, even with a hypothetical future mass spectrometer that could be 10x higher resolution, 10x more sensitive, 10x better dynamic range, 10x less expensive, and 10x easier to use, we may not be any closer to identifying the distinct (and almost by definition less common) analytes that differentiate study conditions if we were to still continue to use informatics algorithms that search only the genomics-derived FASTA space (with only one or two PTMs considered) to then quantify only those identified-in-FASTA-search-space analytes.

For example, in the main paper, we reference a recent and exciting Alzheimer study in which the two proteins that the authors proposed that separated the study conditions were not ones which would have been found in a standard FASTA file without any PTMs considered: instead, the first proposed protein included two extra amino acids at its C-terminal end; and the second proposed protein's distinguishing feature was the presence of a PTM (phosphorylation) at a specific site (vs the unphosphorylated form).

However, even with today's instruments, ~75-95% of analytes in the MS are being unused by existing DIA algorithms, yet they contain, by definition, at least some portion of these unexpected but biologically valuable sequences (e.g., splice variants, sequence variation, protein precursors, unusual proteolytic cleavages, novel proteins, etc.).

In a different but related manner, since PTMs play a disproportionate role in biology, existing algorithms that allow for only one or two modifications to be searched and therefore ignore >99% of other possible PTMs make it far less likely that one would identify peptides that play a disproportionate role in differentiating study conditions. And similarly to the unexpected sequence scenario described above, the ~75-95% of analytes in the MS that current algorithms are not using contain, by definition, at least some portion of these unexplored but potentially valuable PTMs (e.g., glycosylation, citrullination, ubiquitination, acetylation, phosphorylation, methylation, sumoylation, and hundreds more).

Lastly, we asked ChatGPT (using model 4O available from 12/10/2024) the following question:

> *"Would it be fair to say that PTMs play a disproportionate role in disease? Similarly, would it be fair to say that splice variants, sequence variation, protein precursors, unusual proteolytic cleavages, or novel proteins play a disproportionate role in disease? Why or why not? Please cite your sources using the citation format that Nature Method requires."*

And we received the answer below. This answers matches our intuition and our understanding, but we encourage the reader to review the answers by themselves, with other experts, with their own favorite AI tools, or some combination of all three:

**Excerpt 1: Response from ChatGpt (using model 4O from 12/10/2024):**

To address whether post-translational modifications (PTMs), splice variants, sequence variations, protein precursors, unusual proteolytic cleavages, or novel proteins play a disproportionate role in disease, it's essential to evaluate the evidence for their involvement in disease mechanisms and progression. Below is an analysis with citations formatted as required by *Nature Methods*.

---

### Post-Translational Modifications (PTMs)

PTMs indeed play a **disproportionate role in disease** due to their regulatory impact on protein function, localization, and interaction networks. Aberrant PTMs can drive disease states, including cancer, neurodegeneration, and autoimmune disorders. For example:

- **Phosphorylation** dysregulation is critical in cancer, as hyperphosphorylation often activates oncogenes or inactivates tumor suppressors (Hunter, 1995).
- **Glycosylation** abnormalities are implicated in tumor metastasis and immune evasion (Pinho & Reis, 2015).
- **Ubiquitination** and proteasomal degradation defects contribute to neurodegenerative diseases like Parkinson's and Alzheimer's (Goldberg, 2003).

These observations underscore how minor changes in PTM profiles can have system-wide effects, linking them disproportionately to diseases.


---

### Splice Variants

Splice variants can have **disproportionate disease impact** as they frequently generate protein isoforms with altered functions. Alternative splicing can drive tissue-specific expression patterns, but its dysregulation is implicated in diseases such as:

- **Cancer**, where aberrant splicing alters key pathways, such as apoptosis and cell cycle regulation (Oltean & Bates, 2014).

- **Neurological disorders**, where mutations affecting splicing regulators lead to pathological isoforms, as in spinal muscular atrophy (Cieply & Carstens, 2015).


---

### Sequence Variation

Sequence variations, such as **single nucleotide polymorphisms (SNPs)**, can profoundly affect protein function and susceptibility to diseases:

- Variations in the **BRCA1/2** genes increase breast and ovarian cancer risk (Miki et al., 1994).
- SNPs in **APOE** are strongly associated with Alzheimer's disease susceptibility (Corder et al., 1993).

While sequence variations contribute significantly to disease, their role is often one of many contributors in complex disease pathways, making them proportionate rather than disproportionate contributors.


---

### Protein Precursors and Unusual Proteolytic Cleavages

Protein precursors and atypical cleavages often have **disproportionate roles in disease** due to their regulatory functions:

- **Amyloid precursor protein (APP)** cleavage by β-secretase generates amyloid-β peptides, central to Alzheimer's pathology (Haass & Selkoe, 2007).
- Aberrant proteolytic activation of zymogens, such as plasminogen to plasmin, contributes to thrombosis and stroke (Collen, 1999).


---

#### Novel Proteins

Novel proteins, such as those arising from non-canonical translation or cryptic transcriptional start sites, may play unique but not necessarily disproportionate roles. Their function is less well-characterized, but they are increasingly linked to autoimmune diseases and cancer through neoantigen generation (Steinman, 1996).


---

#### Conclusion

PTMs, splice variants, and proteolytic cleavages play **disproportionate roles in disease** due to their systemic and regulatory impacts. Sequence variations and novel proteins are vital contributors but tend to play a more proportional role, as their effects depend on broader genomic and environmental contexts. These distinctions reflect the interconnected nature of molecular biology and disease mechanisms.

In short, ignoring the ~75-95% of analytes in our DIA-MS datasets is deeply unfortunate, particularly considering how precious human samples are – and how important making advancements to human health is.

**Note S5: Query Statements in Google BigQuery (BQ) to Produce Correlation Proofs**

1. From this paper's algorithm Portal, click "Export to CSV" to export quantified analytes to csv file. This will contain all the quantified anlalytes (identified or not)
2. Import csv into BQ as `deepsearchdev.dataset1.ghilresultsv2`
3. Run this query for "Identified Analytes" Correlation For 12.5pg to 500ng inclusive (excluding iRT peptides)

```
--drop table `deepsearchdev.ds.ibd-ghl-correl-for-ided-analytes`;
CREATE TABLE
`deepsearchdev.ds.ibd-ghl-correl-for-ided-analytes` AS
SELECT
id,
```

```
Precursor_Charge,
Modified_Sequence,
CORR(Precursor_Quantity, concentration) AS correlation
FROM (
SELECT
t1.Id_ id,
t1.charge Precursor_Charge,
t1.Peptide_Sequence__Incl_Mods_ Modified_Sequence,
t1.Sample,
t1.Qty__Not_Normalized_ Precursor_Quantity,
t2.concentration
FROM
`deepsearchdev.dataset1.ghilresultsv4` AS t1
INNER JOIN
`deepsearchdev.ds.ibdconc` AS t2
ON
t1.sample = t2.run
-- and now, let's exclude iRT peptides
AND t1.Peptide_Sequence__Incl_Mods_ NOT IN ("LGGNEQVTR",
"YILAGVENSK",
"GTFIIDPGGVIR",
"GTFIIDPAAVIR",
"GAGSSEPVTGLDAK",
"TPVISGGPYEYR",
"VEATFGVDESNAK",
"TPVITGAPYEYR",
"DGLDAASYYAPVR",
"ADVTPADFSEWSK",
"LFLQFGAQGSPFLK" )
-- per Qin's work on DIA-NN, we're focused on peptide concentration from 12.5pg to
500ng (the expected linear range)
AND t2.concentration <= 500
-- only pull, for now, the identified analytes
AND t1.Peptide_Sequence__Incl_Mods_ IS NOT NULL )
GROUP BY
id,
Precursor_Charge,
Modified_Sequence
ORDER BY
Precursor_Charge,
Modified_Sequence;
```

(which is saved as “ibd-ghl-create-correl")

4. Then, run:

```
WITH correlation_counts AS (
SELECT
CASE
```

```
WHEN correlation >= 0.80 AND correlation <= 1.00 THEN '0.80 to 1.00'
--WHEN correlation >= 0.80 AND correlation < 0.90 THEN '0.80 to 0.90'
ELSE '-1 to 0.80'
END AS correlation_bucket
FROM
`deepsearchdev.ds.ibd-ghl-correl-for-ided-analytes`
),
bucket_frequencies AS (
SELECT
correlation_bucket,
COUNT(*) AS frequency
FROM
correlation_counts
GROUP BY
correlation_bucket
)
SELECT
correlation_bucket,
frequency,
ROUND(frequency / SUM(frequency) OVER() * 1, 2) AS frequency_percentage
FROM
bucket_frequencies
ORDER BY
correlation_bucket DESC;
```

```
(which is saved as query "ibd-ghl-histogram")
```

To get:

| correlation_bucket | frequency | frequency_percentage |
|---|---|---|
| 0.80 to 1.00 | 5934 | 93% |
| -1 to 0.80 | 447 | 7% |

### Now, let's compare to DIA-NN:

5. Run this query:

```
--drop table `deepsearchdev.ds.ibd-diann-correl-for-ided-analytes`;

create table `deepsearchdev.ds.ibd-diann-correl-for-ided-analytes` as
SELECT
Precursor_Charge,
Modified_Sequence,
CORR(Precursor_Quantity, concentration) AS correlation
FROM (

SELECT Precursor_Charge,Modified_Sequence,t1.run,Precursor_Quantity,t2.concentration
```

```
FROM `deepsearchdev.ds.ibd-dilution-diann` as t1
inner join `deepsearchdev.ds.ibdconc` as t2
on t1.run=t2.run
-- and now, let's exclude iRT peptides
and t1.Modified_Sequence not in
("LGGNEQVTR","YILAGVENSK","GTFIIDPGGVIR","GTFIIDPAAVIR","GAGSSEPVTGLDAK","TPVISGGPYEYR
","VEATFGVDESNAK","TPVITGAPYEYR","DGLDAASYYAPVR","ADVTPADFSEWSK","LFLQFGAQGSPFLK" )
-- per Qin's work on DIA-NN, we're focused on peptide concentration from 12.5pg to
500ng (the expected linear range)
and t2.concentration <= 500
)
GROUP BY
Precursor_Charge,
Modified_Sequence
ORDER BY
Precursor_Charge,
Modified_Sequence;
```

(which is saved as "ibd-diann-create-histogram")

6. Then, run this query:

```
WITH correlation_counts AS (
SELECT
CASE
WHEN correlation >= 0.80 AND correlation <= 1.00 THEN '0.80 to 1.00'
--WHEN correlation >= 0.80 AND correlation < 0.90 THEN '0.80 to 0.90'
ELSE '-1 to 0.80'
END AS correlation_bucket
FROM
`deepsearchdev.ds.ibd-diann-correl-for-ided-analytes`
),
bucket_frequencies AS (
SELECT
correlation_bucket,
COUNT(*) AS frequency
FROM
correlation_counts
GROUP BY
correlation_bucket
)
SELECT
correlation_bucket,
frequency,
ROUND(frequency / SUM(frequency) OVER() * 1, 2) AS frequency_percentage
FROM
bucket_frequencies
ORDER BY
correlation_bucket DESC;
```

(which is saved as “ibd-diann-histogram")

Which results in this:

| correlation_bucket | frequency | frequency_percentage |
|---|---|---|
| 0.80 to 1.00 | 7752 | 88% |
| -1 to 0.80 | 1104 | 12% |

7. So, from the above analysis (steps 1 through 6 inclusive), we see that: this paper's algorithm’s % of R-Squared >=0.80 is 93%, whereas DIA-NN's (despite its use of a known FASTA library) is 88%.
8. Redoing the above steps but with an Rsquared cutoff of >= 0.90 yields the following result: this paper's algorithm = % and DIA-NN = 75%, as illustrated here:

#### From this paper's algorithm:

| correlation_bucket | frequency | frequency_percentage |
|---|---|---|
| 0.90 to 1.00 | 5292 | 83% |
| -1 to 0.90 | 1089 | 17% |

#### From DIA-NN:

| correlation_bucket | frequency | frequency_percentage |
|---|---|---|
| 0.90 to 1.00 | 6,611 | 75% |
| -1 to 0.90 | 2,245 | 25% |

#### From this paper's algorithm:

| correlation_bucket | frequency | frequency_percentage |
|---|---|---|
| 0.95 to 1.00 | 4300 | 67% |
| -1 to 0.95 | 2081 | 33% |

#### From DIA-NN:

| correlation_bucket | frequency | frequency_percentage |
| --- | --- | --- |
| 0.95 to 1.00 | 5532 | 62% |
| -1 to 0.95 | 3324 | 38% |

9. Now, let's examine the 6 selected peptides that interested Qin using this query:

```
SELECT
Modified_Sequence,
Precursor_Charge,
MAX(correlation) AS bestCorrel
FROM (
SELECT
id,
Precursor_Charge,
Modified_Sequence,
CORR(Precursor_Quantity, concentration) AS correlation
FROM (
SELECT
t1.Id_ id,
t1.charge Precursor_Charge,
t1.Peptide_Sequence__Incl_Mods_ Modified_Sequence,
t1.Sample,
t1.Qty__Not_Normalized_ Precursor_Quantity,
t2.concentration
FROM
`deepsearchdev.dataset1.ghilresultsv4` AS t1
INNER JOIN
`deepsearchdev.ds.ibdconc` AS t2
ON
t1.sample = t2.run
-- and now, let's exclude iRT peptides
AND t1.Peptide_Sequence__Incl_Mods_ IN ("ESDTSYVSLK",
"SDVVYTDWK",
"ILEGFQPSGR",
"YVGGQEHFAHLLILR",
"IADVTSGLIGGEDGR",
"C(Carbamidomethyl)EAC(Carbamidomethyl)PPGYSGPTHQGVGLAFAK")
-- per Qin's work on DIA-NN, we're focused on peptide concentration from 12.5pg to
500ng (the expected linear range)
AND t2.concentration <= 500
-- only pull, for now, the identified analytes
AND t1.Peptide_Sequence__Incl_Mods_ IS NOT NULL
-- and finally, only pull in high quality identifications
--and t1.ReductionFactor = 1
--and t1.__A__MS1_2_m_zs >= 11
)
GROUP BY
id,
Precursor_Charge,
Modified_Sequence )
GROUP BY
Precursor_Charge,
```

```
Modified_Sequence
ORDER BY
Modified_Sequence,
Precursor_Charge
```

With saved query name of “ibd-ghl-qin-5-pep" and which resulted in this:

| Modified_Sequence | Charge | Correlation |
|---|---|---|
| C(Carbamidomethyl)EAC(Carbamidomethyl)PPGYSGPTHQGVGLAFAK | 3 | 0.98 |
| ESDTSYVSLK | 2 | 0.97 |
| IADVTSGLIGGEDGR | 2 | 0.97 |
| ILEGFQPSGR | 2 | 0.98 |
| SDVVYTDWK | 2 | 0.99 |
| YVGGQEHFAHLLILR | 2 | 1.00 |
| YVGGQEHFAHLLILR | 3 | 0.99 |
| YVGGQEHFAHLLILR | 4 | 0.99 |

To plot each peptide, run query below and the export last 2 columns to excel, and then plot within excel....

```
SELECT
t1.Id_,
t1.Peptide_Sequence__Incl_Mods_ Modified_Sequence,
t1.charge Precursor_Charge,
t1.Sample,
t2.concentration as Concentration_ng,
t1.Qty__Not_Normalized_ Relative_Quantity_cps
FROM
`deepsearchdev.dataset1.ghilresultsv4` AS t1
INNER JOIN
`deepsearchdev.ds.ibdconc` AS t2
ON
t1.sample = t2.run
-- and now, let's exclude iRT peptides
and t1.Charge=4
AND t1.Peptide_Sequence__Incl_Mods_ =
--"ESDTSYVSLK"
--"SDVVYTDWK"
-- "ILEGFQPSGR"
"YVGGQEHFAHLLILR"
--"IADVTSGLIGGEDGR"
--"C(Carbamidomethyl)EAC(Carbamidomethyl)PPGYSGPTHQGVGLAFAK"
--)
-- per Qin's work on DIA-NN, we're focused on peptide concentration from 12.5pg to
500ng (the expected linear range)
AND t2.concentration <= 500
```

```
AND t1.Qty__Not_Normalized_ IS NOT NULL
-- only pull, for now, the identified analytes
AND t1.Peptide_Sequence__Incl_Mods_ IS NOT NULL
-- and finally, only pull in high quality identifications
--and t1.ReductionFactor = 1
--and t1.__A__MS1_2_m_zs >= 11
ORDER BY
t1.Id_,
modified_sequence,
precursor_charge,
concentration,
sample
```

For DIA-NN, run this:

```
SELECT
Modified_Sequence, Precursor_Charge, correlation
FROM
`deepsearchdev.ds.ibd-diann-correl-for-ided-analytes`
where Modified_Sequence in (
"ESDTSYVSLK",
"SDVVYTDWK",
"ILEGFQPSGR",
"YVGGQEHFAHLLILR",
"IADVTSGLIGGEDGR",
"C(UniMod:4)EAC(UniMod:4)PPGYSGPTHQGVGLAFAK"
)
order by Modified_Sequence, Precursor_Charge
```

To get this:

| Modified_Sequence | Precursor_Charge | correlation |
|---|---|---|
| C(UniMod:4)EAC(UniMod:4)PPGYSGPTHQGVGLAFAK | 3 | 0.99 |
| ESDTSYVSLK | 2 | 0.99 |
| IADVTSGLIGGEDGR | 2 | 0.99 |
| ILEGFQPSGR | 2 | 1.00 |
| SDVVYTDWK | 2 | 0.99 |
| YVGGQEHFAHLLILR | 2 | 1.00 |
| YVGGQEHFAHLLILR | 3 | 0.99 |
| YVGGQEHFAHLLILR | 4 | 0.99 |
| | average | 0.99 |

================================

10. Now, let's consider the quantitative accuracy for non-identified analytes. We ran this query:

```
DROP TABLE `deepsearchdev.ds.ibd-ghl-correl-for-non-id-high-quality-analytes`;

CREATE TABLE
`deepsearchdev.ds.ibd-ghl-correl-for-non-id-high-quality-analytes` AS
SELECT
id,
Precursor_Charge,
Modified_Sequence,
CORR(Precursor_Quantity, concentration) AS correlation
FROM (
SELECT
t1.Id_ id,
t1.charge Precursor_Charge,
t1.Peptide_Sequence__Incl_Mods_ Modified_Sequence,
t1.Sample,
t1.Qty__Not_Normalized_ Precursor_Quantity,
t2.concentration
FROM
`deepsearchdev.dataset1.ghilresultsv4` AS t1
INNER JOIN
`deepsearchdev.ds.ibdconc` AS t2
ON
t1.sample = t2.run
-- per Qin's work on DIA-NN, we're focused on peptide concentration from 12.5pg to
500ng (the expected linear range)
AND t2.concentration <= 500
-- only pull, for now, the identified analytes
AND t1.Peptide_Sequence__Incl_Mods_ IS NULL
-- and finally, only pull in high quality identifications
AND t1.ReductionFactor = 1
AND t1.__A__MS1_2_m_zs >= 0 )
GROUP BY
id,
Precursor_Charge,
Modified_Sequence
ORDER BY
Precursor_Charge,
Modified_Sequence;
```

And then:

```
WITH
correlation_counts AS (
SELECT
CASE
WHEN correlation >= 0.90 AND correlation <= 1.00 THEN '0.90 to 1.00'
--WHEN correlation >= 0.80 AND correlation < 0.90 THEN '0.80 to 0.90'
ELSE '-1 to 0.90'
```

```
END
AS correlation_bucket
FROM
`deepsearchdev.ds.ibd-ghl-correl-for-non-id-high-quality-analytes` ),
bucket_frequencies AS (
SELECT
correlation_bucket,
COUNT(*) AS frequency
FROM
correlation_counts
GROUP BY
correlation_bucket )
SELECT
correlation_bucket,
frequency,
ROUND(frequency / SUM(frequency) OVER() * 1, 2) AS frequency_percentage
FROM
bucket_frequencies
ORDER BY
correlation_bucket DESC;
```

To get this for "high quality non-identified analytes"

| correlation_bucket | frequency | frequency_percentage |
|---|---|---|
| 0.90 to 1.00 | 33890 | 80% |
| -1 to 0.90 | 8285 | 20% |

Now, let's look at mid-quality non-identified analytes:

```
DROP TABLE
`deepsearchdev.ds.ibd-ghl-correl-for-non-id-mid-quality-analytes`;
CREATE TABLE
`deepsearchdev.ds.ibd-ghl-correl-for-non-id-mid-quality-analytes` AS
SELECT
id,
Precursor_Charge,
Modified_Sequence,
CORR(Precursor_Quantity, concentration) AS correlation
FROM (
SELECT
t1.Id_ id,
t1.charge Precursor_Charge,
t1.Peptide_Sequence__Incl_Mods_ Modified_Sequence,
t1.Sample,
t1.Qty__Not_Normalized_ Precursor_Quantity,
t2.concentration
FROM
```

```
`deepsearchdev.dataset1.ghilresultsv4` AS t1
INNER JOIN
`deepsearchdev.ds.ibdconc` AS t2
ON
t1.sample = t2.run
-- per Qin's work on DIA-NN, we're focused on peptide concentration from 12.5pg to
500ng (the expected linear range)
AND t2.concentration <= 500
-- only pull, for now, the identified analytes
AND t1.Peptide_Sequence__Incl_Mods_ IS NULL
-- and finally, only pull in high quality identifications
AND t1.ReductionFactor < 1
AND t1.__A__MS1_2_m_zs >= 11 )
GROUP BY
id,
Precursor_Charge,
Modified_Sequence
ORDER BY
Precursor_Charge,
Modified_Sequence;
```

And

```
WITH
correlation_counts AS (
SELECT
CASE
WHEN correlation >= 0.90 AND correlation <= 1.00 THEN '0.90 to 1.00'
--WHEN correlation >= 0.80 AND correlation < 0.90 THEN '0.80 to 0.90'
ELSE '-1 to 0.90'
END
AS correlation_bucket
FROM
`deepsearchdev.ds.ibd-ghl-correl-for-non-id-mid-quality-analytes` ),
bucket_frequencies AS (
SELECT
correlation_bucket,
COUNT(*) AS frequency
FROM
correlation_counts
GROUP BY
correlation_bucket )
SELECT
correlation_bucket,
frequency,
ROUND(frequency / SUM(frequency) OVER() * 1, 2) AS frequency_percentage
FROM
bucket_frequencies
ORDER BY
correlation_bucket DESC;
```

Which gives us this for the “mid quality non-identified analytes”:

| correlation_bucket | frequency | frequency_percentage |
|---|---|---|
| 0.90 to 1.00 | 27708 | 48% |
| -1 to 0.90 | 29771 | 52% |

Finally, for “low quality non-identified analytes”, we do this:

```
drop table `deepsearchdev.ds.ibd-ghl-correl-for-non-id-low-quality-analytes`;

CREATE TABLE
`deepsearchdev.ds.ibd-ghl-correl-for-non-id-low-quality-analytes` AS
SELECT
id,
Precursor_Charge,
Modified_Sequence,
CORR(Precursor_Quantity, concentration) AS correlation
FROM (
SELECT
t1.Id_ id,
t1.charge Precursor_Charge,
t1.Peptide_Sequence__Incl_Mods_ Modified_Sequence,
t1.Sample,
t1.Qty__Not_Normalized_ Precursor_Quantity,
t2.concentration
FROM
`deepsearchdev.dataset1.ghilresultsv4` AS t1
INNER JOIN
`deepsearchdev.ds.ibdconc` AS t2
ON
t1.sample = t2.run
-- per Qin's work on DIA-NN, we're focused on peptide concentration from 12.5pg to
500ng (the expected linear range)
AND t2.concentration <= 500
-- only pull, for now, the identified analytes
AND t1.Peptide_Sequence__Incl_Mods_ IS NULL
-- and finally, only pull in high quality identifications
AND t1.ReductionFactor < 1
AND t1.__A__MS1_2_m_zs < 11 )
GROUP BY
id,
Precursor_Charge,
Modified_Sequence
ORDER BY
Precursor_Charge,
Modified_Sequence;
```

And

```
WITH
correlation_counts AS (
SELECT
CASE
WHEN correlation >= 0.90 AND correlation <= 1.00 THEN '0.90 to 1.00'
--WHEN correlation >= 0.80 AND correlation < 0.90 THEN '0.80 to 0.90'
ELSE '-1 to 0.90'
END
AS correlation_bucket
FROM
`deepsearchdev.ds.ibd-ghl-correl-for-non-id-low-quality-analytes` ),
bucket_frequencies AS (
SELECT
correlation_bucket,
COUNT(*) AS frequency
FROM
correlation_counts
GROUP BY
correlation_bucket )
SELECT
correlation_bucket,
frequency,
ROUND(frequency / SUM(frequency) OVER() * 1, 2) AS frequency_percentage
FROM
bucket_frequencies
ORDER BY
correlation_bucket DESC;
```

To give us:

| | | |
|---|---|---|
| 0.90 to 1.00 | 4014 | 23% |
| -1 to 0.90 | 13471 | 77% |


====================== END OF SUPPLEMENTAL ======================
